# Choice suppression is achieved through opponent but not independent function of the striatal indirect pathway in mice

**DOI:** 10.1101/675850

**Authors:** Kristen Delevich, Benjamin Hoshal, Anne GE Collins, Linda Wilbrecht

## Abstract

The dorsomedial striatum (DMS) plays a key role in action selection, but little is known about how direct and indirect pathway spiny projection neurons (dSPNs and iSPNs) contribute to choice suppression in freely moving animals. Here, we used pathway-specific chemogenetic manipulation during a serial choice foraging task to test opposing predictions for iSPN function generated by two theories: 1) the ‘select/suppress’ heuristic which suggests iSPN activity is required to suppress alternate choices and 2) the network-inspired <u>Op</u>ponent <u>A</u>ctor <u>L</u>earning model (OpAL) which proposes that the weighted difference of dSPN and iSPN activity determines choice. We found that chemogenetic activation, but not inhibition, of iSPNs disrupted learned suppression of nonrewarded choices, consistent with the predictions of the OpAL model. Our findings suggest that iSPNs’ role in stopping and freezing does not extend in a simple fashion to choice suppression. These data may provide insights critical for the successful design of interventions for addiction or other conditions in which suppression of behavior is desirable.

## Introduction

In everyday decision-making, we often consider multiple options in a serial fashion, foregoing low value choices in order to arrive at a higher value choice. As time passes and the environment or our needs change, we also learn to suppress formerly high value choices to adjust our behavior to new contingencies or states. The inability to suppress problematic choices is a core component of addiction, eating disorders, and obsessive-compulsive disorder (Lucantonio et al., 2012; Gillan et al., 2015; Kessler et al., 2016). The mechanisms underlying learned choice suppression are therefore highly relevant to psychiatry and public health.

The DMS (homologous to the primate caudate) is a key brain structure for goal-directed action selection (Yin et al., 2005a; Yin et al., 2005b; Balleine et al., 2007; Balleine and O’Doherty, 2010; Shan et al., 2014), and striatal dysfunction is associated with maladaptive choice behavior (Everitt et al., 2008; Castane et al., 2010; Foerde et al., 2015; Volkow and Morales, 2015; Friedman et al., 2017). However, it is still not well understood how choice selection and suppression are implemented at the circuit level (Cox and Witten, 2019). Furthermore, much of the relevant functional data comes from two-alternative forced choice (2AFC) tasks (Hikosaka et al., 2006; Lau and Glimcher, 2008; Kim et al., 2009; Tai et al., 2012; Donahue et al., 2018), in which movement is constrained and it is difficult to dissociate the selection of one choice (e.g. turn left) from the suppression of another (e.g. do not turn right). Therefore, studying DMS function in the context of a serial task in which animals move freely and select among multiple options may reveal new insights into the circuit mechanisms that underlie choice selection and suppression. These findings may also be more relevant to human contexts in which choice is often performed in a serial fashion and is not tightly tied to motor space or motor inhibition, for example choosing among multiple activities available in a kitchen (eating, drinking, cooking, cleaning, etc.) or choosing among multiple options (to drink water, soda, beer or wine?).

The DMS is primarily composed of dopamine D1 receptor expressing dSPNs and dopamine D2 receptor expressing iSPNs (Gerfen et al., 1990), whose activity reflect task features including movement, cues, and value (Samejima et al., 2005; Lau and Glimcher, 2008; Thorn et al., 2010; Seo et al., 2012; Isomura et al., 2013; Nonomura et al., 2018; Shin et al., 2018). Consistent with predictions from functional neuroanatomy (Albin et al., 1989; Alexander and Crutcher, 1990; Mink, 1996) and theoretical work (Frank et al., 2004; Collins and Frank, 2014), optogenetic stimulation of dSPNs promotes movement and reinforces actions (‘go’ functions) (Kravitz et al., 2010; Kravitz et al., 2012; Yttri and Dudman, 2016) whereas optogenetic stimulation of iSPNs inhibits movement and drives aversion (‘no go’ functions) (Kravitz et al., 2010; Kravitz et al., 2012; Yttri and Dudman, 2016). In a 2AFC task, dSPN stimulation promotes contraversive choices whereas iSPN stimulation promotes ipsiversive choices in a manner that is reward history dependent (Tai et al., 2012). While these data suggest that the function of dSPNs and iSPNs in decision-making are dichotomous, *in vivo* recordings have shown that they are co-active during goal-directed movement (Cui et al., 2013; Isomura et al., 2013; Jin et al., 2014). To reconcile these observations, it has been proposed that the two pathways work in concert such that dSPNs promote desired actions/choices whereas iSPNs suppress competing actions/choices (Alexander and Crutcher, 1990; Mink, 1996; Hikosaka et al., 2000; Cui et al., 2013), but this interpretation, which we refer to as the ‘select/suppress’ heuristic, has not been tested in the context of serial choice.

In the current study, we compared the predictions generated by the ‘select/suppress’ heuristic and a network-inspired algorithmic model known as the <u>Op</u>ponent <u>A</u>ctor <u>L</u>earning model (OpAL) in which the weighted difference in dSPN and iSPN activity determines choice. We found that the two related models make opposite predictions about how iSPN manipulation should affect learned choice suppression. We then tested the model predictions by performing chemogenetic manipulation of iSPNs and dSPNs in mice trained in a freely moving serial choice task. We found that excitation, but not inhibition of iSPNs disrupted learned choice suppression in this context, consistent with the OpAL model predictions.

## Results

In order to quantify selection and suppression of choices, we trained mice in an odor-guided serial choice task in which mice approach multiple distinctly scented pots in a serial fashion, rejecting pots until they choose one by digging in the scented shavings it contains (Johnson et al., 2016) (Fig. 1a). Only one odor was rewarded (O1, anise), and the odor-action-reward contingency was learned through trial and error during an Acquisition Phase. Training-naïve mice consistently exhibit preference for one nonrewarded odor (O2, thyme). Therefore, in addition to learning to choose O1, a large part of the Acquisition Phase involved learning to suppress choice to O2 (Fig. 1b-c). Twenty-four hours later mice entered a recall Test Phase where their ability to select the rewarded odor (O1) and suppress choice to the remaining nonrewarded odors was assessed (Fig. 1a,c and Methods).

**Fig. 1.**
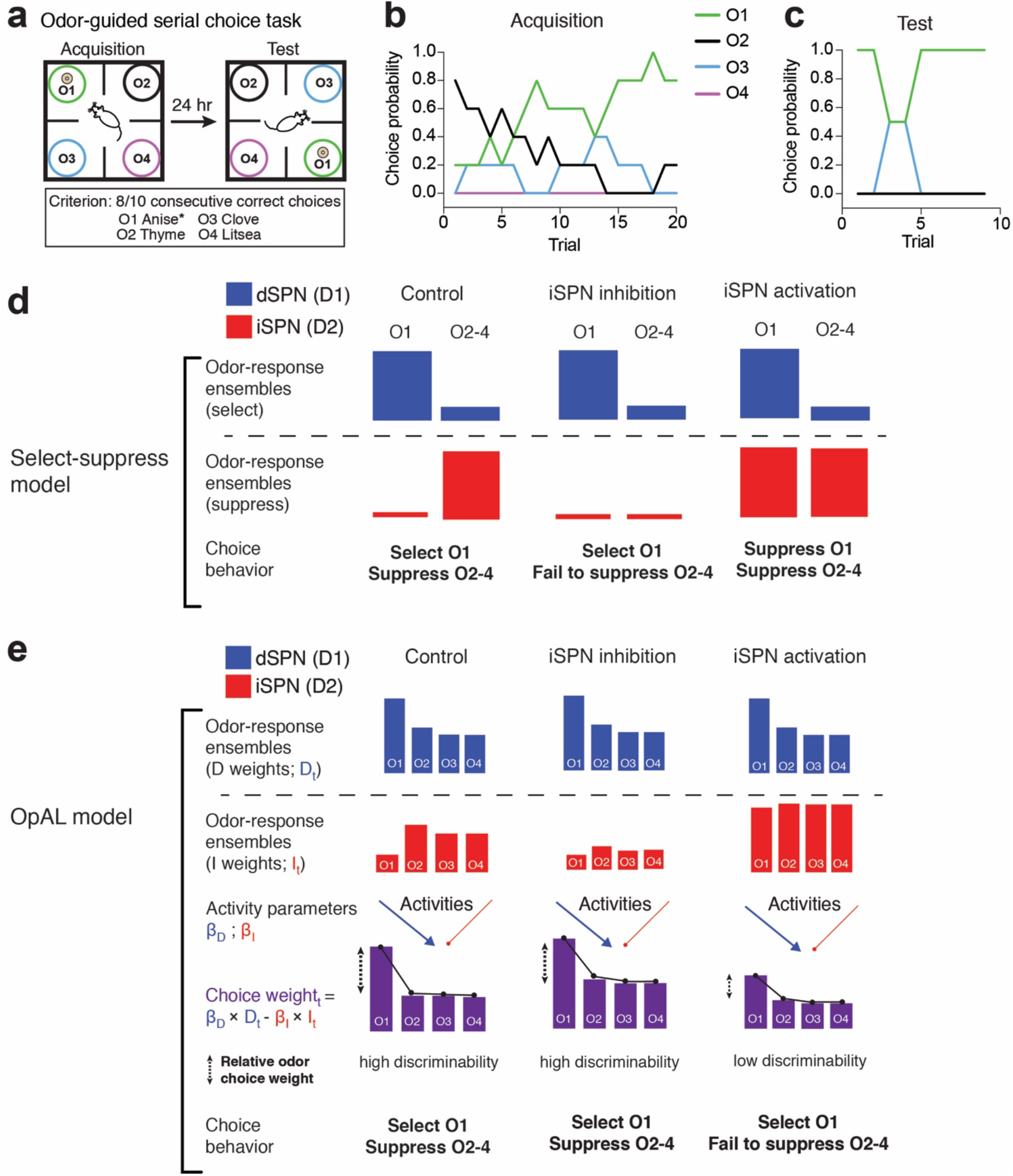
The ‘select/suppress’ heuristic and OpAL model make different predictions for iSPN manipulation effects on odor-guided serial choice task performance. (**a**) 4 option odor-based serial choice task. **(b)** Odor choices made over the Acquisition session in a representative mouse. Training-naïve mice consistently exhibit preference for the nonrewarded odor 2 (O2, thyme) and learn through trial and error that Odor 1 (O1, anise) is rewarded until they reach a criterion of 8/10 consecutive choices to O1. **(c)** Odor choices made during the Test Phase 24 hours later by the same mouse, showing successful suppression of training-naive preferred O2 choices. In **(b)** and **(c)**, choice probability was calculated over a two-trial rolling average. (**d**) A ‘select/suppress’ model emphasizes the independent role of iSPNs in suppressing choices to nonrewarded odor options and predicts that iSPN inhibition will cause failure to suppress low value choices whereas iSPN activation will enhance suppression of low value choices or increase omissions. **(e)** The <u>Op</u>ponent <u>A</u>ctor <u>L</u>earning model. Here, weights onto dSPNs and iSPNs for different odor stimuli reflect learned values updated by a trial and error RL mechanism during the Acquisition Phase. A unique feature of OpAL is that dSPN and iSPN value updates emphasize positive and negative prediction errors, respectively. Weights are scaled by separate activity parameters, *β*_*D*_ and *β*_*I*_ that modulate the extent to which dSPN and iSPN pathways influence choice (this scaling can capture modulation e.g. by chemogenetics). Finally, the weighted difference between dSPN activity and iSPN (Choice weights) are transformed into choice probabilities using the softmax function. OpAL predicts that 1) inhibition of iSPNs increases choice weights across odor options but that the relative value difference between odor choice weights is maintained and 2) activation of iSPN minimizes the difference in choice weights across odor choices which would lower value discriminability among odor choice weights and render choice behavior more stochastic. Increased choices to nonrewarded odors is indicative of a failure to suppress choices that the animal has learned are low in value.

### ‘Select/suppress’ and OpAL models make different predictions for how iSPN activity relates to choice suppression

If iSPNs are responsible for choice suppression as suggested by the ‘select/suppress’ heuristic (Fig. 1d), then inhibition of iSPNs during Test Phase should lead to more choices to Odors 2-4, indicating a failure to suppress choice to nonrewarded odors (Fig. 1d). Conversely, activation of iSPNs should facilitate suppression of nonrewarded choices and improve performance, or alternatively, produce choice omissions (Fig. 1d). Importantly, the select/suppress model is a heuristic that doesn’t provide a computational explanation for how dSPN and iSPN activity affects choice behavior.

Alternate, network-inspired models of basal ganglia function (Collins and Frank, 2014) and recent *in vivo* data suggest that an independent division of labor does not properly account for how action and choice arise from the activity of dSPN and iSPN populations (Barbera et al., 2016; Klaus et al., 2017; Markowitz et al., 2018; Meng et al., 2018; Parker et al., 2018; Shin et al., 2018). A model that accounts for the opponent nature of the two pathways may generate different and more accurate predictions. Therefore, we turned to OpAL, an algorithmic model of a biologically plausible basal ganglia network (Fig. 1e and S1) in which weights onto dSPN and iSPN ensembles representing a given odor option are updated in a manner that emphasizes the positive prediction errors in dSPNs (D weights) and negative prediction errors in iSPNs (I weights). These D and I weights are scaled independently to reflect *in vivo* dSPN and iSPN population activity, and choice weights controlling the decision policy are computed as the difference between dSPN and iSPN population activity for each available odor (Fig. 1f and S1, see Methods for more details), reflecting their downstream competitive output. OpAL predicted 1) that inhibiting iSPN activity would not alter the relative difference between choice weights, leaving value discriminability among odors options high (Fig. 1f), and 2) that activating iSPNs would reduce the difference in choice weights between the rewarded odor and the nonrewarded odors, lowering discriminability and making choice behavior more stochastic (Fig. 1f and S1).

We next simulated performance in the odor-guided serial choice task by adjusting iSPN population activity during Test Phase via parameters that mimicked chemogenetic activation or inhibition (Fig. 2a). To capture the preference that training-naïve mice exhibit for O2, the D weight for O2 was initialized at 1.5 and 1 for all other odors (see Methods for more detail). During the Acquisition Phase, when no manipulation is applied, all groups took a similar number of choices to reach criterion (ANOVA main effect p>0.99; Fig. 2b). OpAL simulations predicted that activation of iSPNs would increase Test Phase choices to criterion while inhibition of iSPNs should not affect performance (ANOVA main effect ***p<0.0001); Fig. 2c). Furthermore, OpAL simulations predicted that iSPN activation would lead to significantly more choices of nonrewarded odors, especially O2, suggesting a reduction in ability to suppress choice based on learned information. In sum, OpAL made predictions regarding iSPN function that were opposite to those generated by the ‘select/suppress’ heuristic.

**Fig. 2.**
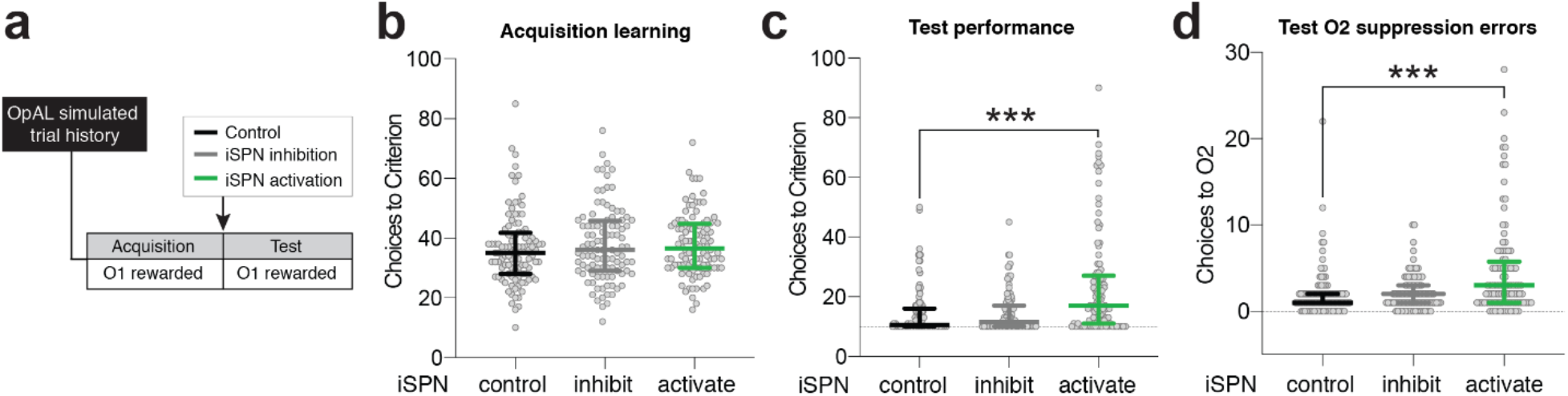
OpAL model predicted that iSPN activation, but not inhibition will impair learned suppression of nonrewarded choices. (**a**) OpAL was used to simulate trial histories (N=100) for the odor-guided serial choice task under three conditions: 1) control and with DREADD manipulation to 2) ‘inhibit’ or 3) ‘activate’ iSPNs during the Test Phase of the task. The OpAL simulation predicted that **(b)** choices to criterion during Acquisition Phase (when no treatment was applied) would not differ across the three conditions (p= 0.45); **(c)** iSPN activation during Test Phase would significantly increase choices to criterion compared to controls (***p<0.0001), while inhibition would not significantly affect performance (p>0.99); and **(d)** iSPN activation during Test Phase would significantly increase O2 choices, indicating failure to suppress choice of the training-naive high value option that mice had learned to avoid (***p<0.0001). Data are presented as median ± IQR. Hypothesis tests were conducted using Kruskal-Wallis ANOVA with Dunn’s test for multiple comparisons.

### Chemogenetic activation, not inhibition, of the indirect pathway impairs the ability to suppress nonrewarded choices

To directly test the predictions made by the ‘select/suppress’ heuristic and OpAL model we turned to *in vivo* chemogenetic manipulation. First, the efficacy of activating and inhibitory DREADD manipulation was confirmed in slice electrophysiology experiments; CNO activation of hM4Di suppressed iSPN synaptic release (Fig. 3a-f) and CNO activation of hM3Dq depolarized iSPNs (Fig. 3g,h). There was a low incidence of colocalization of mCherry and choline acetytransferase (ChAT) in D2-Cre mice (5/146 ChAT+ neurons located within regions of mCherry expression), indicating that infection was largely restricted to iSPNs (Fig. 3i,j).

**Fig. 3.**
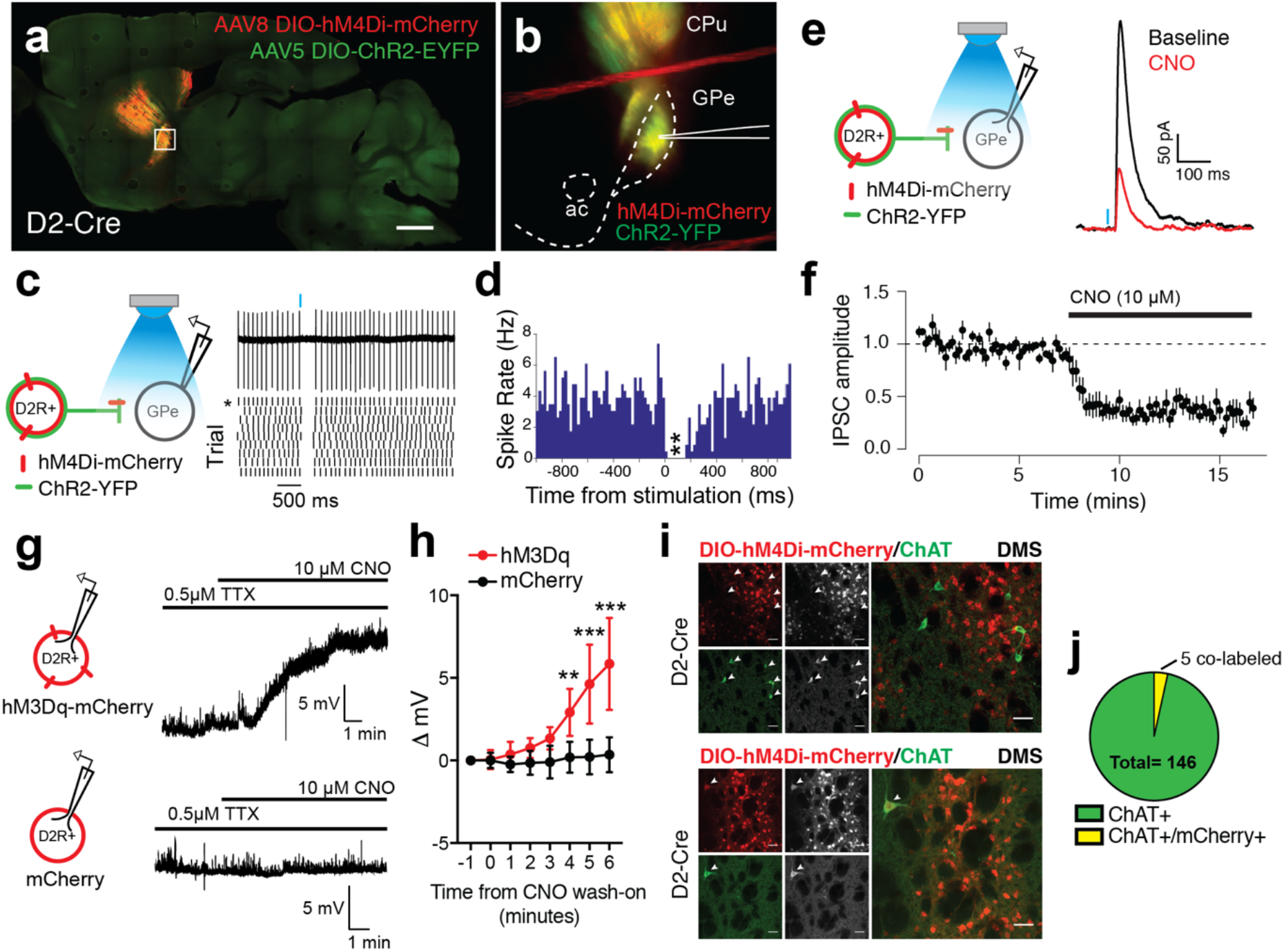
Selective modulation of dorsomedial striatum (DMS) SPNs using designer receptor exclusively activated by designer drugs (DREADDs). (**a)** DMS of D2-Cre were transduced with a 2:1 mixture of Cre-dependent hM4Di-mCherry and ChR2-EYFP, leading to expression in the indirect pathway. Scale bars indicate 1 mm. **(b)** Acute 300 μM sagittal slices containing GPe were prepared and GPe neurons were targeted for patch-clamp recording. **(c)** Left panel: cell-attached recording configuration; right panel: brief light stimulation (10 ms) causes long-lasting suppression of postsynaptic spiking in example tonically firing GPe neuron. Asterisk indicates raster that corresponds to raw trace above. **(d)** Peristimulus spike histogram indicates that 10 ms light stimulation significantly reduces spontaneous firing rate for 160 ms (>3 z-scores below pre-stimulus mean firing rate; n= 6 cells). **(e)** Left panel: whole-cell recording configuration; right panel: example recording of eIPSC before and after CNO (10μM) wash on. **(f)** Normalized eIPSC amplitude before and after CNO wash on (mean ± SEM plotted for 6 cells). Pre-CNO vs. post-CNO normalized eIPSC amplitude (D=0.9839, ***p<0.0001 two-sample Kolmogorov-Smirnov test). **(g)** D2-Cre mice were injected into DMS bilaterally with DIO-hM3Dq-mCherry virus, and in separate experiments, mCherry+ neurons were patched in DMS of D2-Cre; DIO-hM3Dq-mCherry transfected mice or D2-Cre; DIO-mCherry transfected mice. After establishing stable baseline in current clamp, 10 μM CNO was bath applied in the presence of TTX (0.5 μM). **(h)** CNO application significantly depolarized hM3Dq-mCherry+ but not control mCherry+ cells, beginning in the 3-4 minute time bin after CNO hit bath. There was a significant interaction between virus and time on membrane potential (F(7,63)= 15.64, ***p<0.0001 repeated measures ANOVA). Mean ± 95% confidence interval are plotted (n= 6 hM3Dq, 5 mCherry control neurons). **(i)** Top panel: Representative images showing absence of colocalization of Cre-dependent mCherry (red) and choline acetyltransferase immunoreactivity (ChAT; green) within the DMS of AAV8-hSyn-DIO-mCherry transduced D2-Cre mice (N=4). White arrows indicate location of ChAT+ neurons. Bottom panel: Representative images demonstrating rare colocalization of Cre-dependent mCherry (red) and choline acetyltransferase immunoreactivity (ChAT; green) within the DMS of AAV8-hSyn-DIO-mCherry transduced D2-Cre mice. White arrow indicates location ChAT+ neuron that also exhibits mCherry expression. **(j)** Quantified proportion of ChAT+/mCherry+ neurons identified within regions of AAV8-hSyn-DIO-mCherry infection: 5 out of 146 ChAT+ neurons colocalized with mCherry, corresponding to ~3% colabeling. Scale bars = 50 μM.

To test the role of iSPNs in choice behavior, D2-Cre mice were injected bilaterally into the DMS with 0.5 μL of Cre-dependent DREADD virus (hM4Di-mCherry or hM3Dq-mCherry) and were trained 4-6 weeks later in the 4 option odor-guided serial choice task (Fig. 4a). Mice that expressed Cre-inducible mCherry were used to control for any effects of surgery, AAV infection, and CNO administration on behavior. Prior to Acquisition all mice received i.p. injections of saline (Fig. 4a). No difference in Acquisition learning, measured as choices to criterion was observed across groups (ANOVA main effect p= 0.54) (Fig. 4b). During Acquisition, there was a significant effect of odor identity on nonrewarded choices, with O2 being the most frequently chosen nonrewarded odor (ANOVA main effect p<0.001; Fig. 4c), consistent with previous studies in our lab that found that training-naive mice consistently exhibit a preference for O2 (thyme) (Johnson and Wilbrecht, 2011; Johnson et al., 2016). Twenty-four hours after Acquisition, mice were administered CNO (1.0 mg/kg, i.p.) and run in the Test Phase. mCherry control and D2-Cre inhibitory DREADD (hM4Di) groups exhibited robust recall of the rewarded choice and successful suppression of the nonrewarded choices during Test, with most mice reaching criterion in the minimum number of trials required (10) (Fig. 4d). Mice expressing activating DREADD in iSPNs (D2-hM3Dq) required significantly more choices to reach criterion (ANOVA main effect **p<0.01; Fig. 4d), made more nonrewarded choices (ANOVA main effect *p<0.05; Fig. 4e), and made significantly more choices to O2 compared to mCherry controls (ANOVA main effect *p<0.05; Fig. 4f). Given that O2 was the preferred odor prior to Acquisition learning, these results suggest that activating iSPNs impaired learned suppression. Finally, D2-hM3Dq mice were slower to accumulate rewards during the Test Phase (**p<0.01; Fig. 4g) compared to mCherry controls, whereas D2-hM4Di mice did not significantly differ from mCherry controls (Fig. S2). These data were consistent with the predictions made by the OpAL model (Fig. 1f) but inconsistent with those made by the ‘select/suppress’ model.

**Fig. 4.**
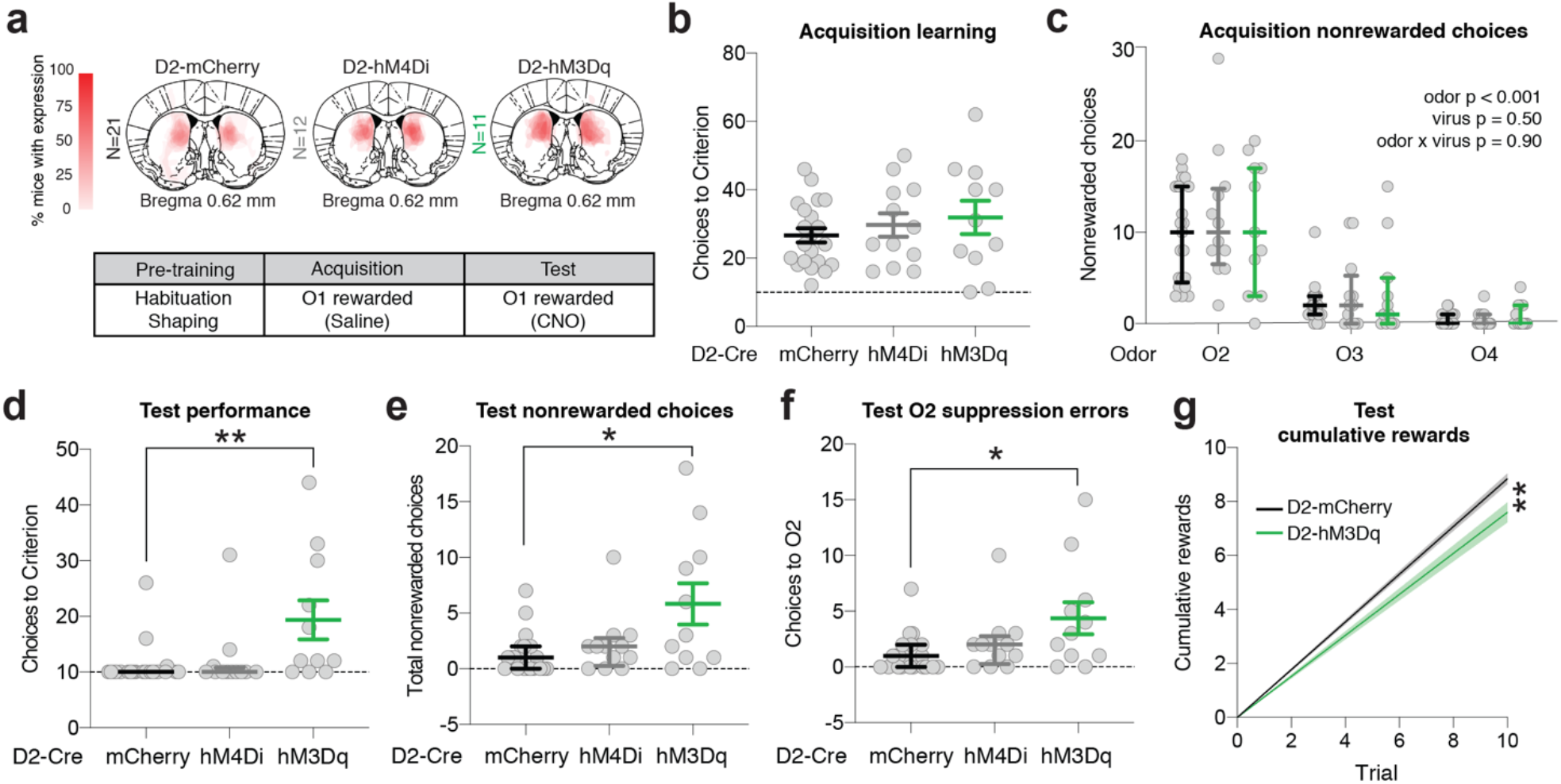
iSPN chemogenetic activation not inhibition impairs suppression of nonrewarded choices. (**a**) Top panel: schematic adapted from (Franklin and Paxinos, 1997) illustrating injection site and viral spread D2-Cre DIO-mCherry (N=21), DIO-hM4Di (N=12), and DIO-hM3Dq (N= 11) mice. Opacity indicates the number of mice in each cohort that expressed virus in a given location. Bottom panel: summary of behavior; during pre-training mice are habituated to the arena and cheerio rewards (habituation) and learn to dig in unscented wood shavings to retrieve buried cheerio reward (shaping); during Acquisition, mice learn that only Odor 1 is rewarded (criterion = 8/10 correct consecutive choices); 24 hours later during Test, mice are tested for their recall of Acquisition learning (criterion = 8/10 correct consecutive choices). Mice received i.p. injection of Saline or 1 mg/kg CNO (clozapine-N-oxide) 30 minutes prior to testing. (**b**) Acquisition (Saline) choices to criterion did not differ across groups (F(2,19.4)=0.64, p=0.54; Welch’s ANOVA). **(c)** There was a significant effect of odor identity (F(1.41, 57.9)= 75.6, **p<0.001 two-way ANOVA) but not virus (F(2, 41)= 0.70, p=0.50 two-way ANOVA) on nonrewarded choices during Acquisition. This shows all 3 groups chose O2 (thyme) more than the other nonrewarded odors, but then suppressed this choice eventually to reach criterion during Acquisition. **(d)** During Test (CNO), D2-hM3Dq mice took significantly more choices to reach criterion compared to mCherry controls (**p<0.01 Kruskal-Wallis ANOVA with Dunn’s test for multiple comparisons). **(e)** D2-hM3Dq mice made more nonrewarded choices compared to mCherry controls (*p<0.05 Kruskal-Wallis ANOVA with Dunn’s test for multiple comparisons). **(f)** D2-hM3Dq mice chose the training-naïve preferred odor (O2) significantly more often during Test, suggesting failure of learned suppression (**p<0.01 Kruskal-Wallis ANOVA with Dunn’s test for multiple comparisons). **(g)** D2-hM3Dq mice were significantly slower to accumulate rewards during the Test Phase compared to mCherry controls (**p<0.01 Kruskal-Wallis ANOVA with Dunn’s test for multiple comparisons). Mean ± SEM are plotted for normally distributed data including all groups in b, and the D2-hM3Dq group in d-g. Otherwise data plotted indicate Median ± IQR. Data in f indicate linear regression line with bands plotting the 95% confidence interval.

### Chemogenetic activation of the direct pathway also impairs ability to suppress nonrewarded choices

We found that activation but not inhibition of iSPNs affected Test Phase performance in the odor-guided foraging task. We next tested whether inhibition of dSPNs affected Test Phase performance which would suggest dissociation between iSPN and dSPN contribution to choice behavior. In the case of dSPNs, the ‘select/suppress’ and OpAL models similarly predict that dSPN inhibition would impair Test Phase performance (Fig. S1). D1-Cre mice were bilaterally transduced with Cre-dependent hM4Di-mCherry control mCherry virus and were trained 4-6 weeks later in the 4 option odor-guided serial choice task (Fig. 5a). Acquisition learning was consistent between groups (Fig. 5b). Choices of nonrewarded odors during Acquisition were comparable between D1-hM4Di and mCherry control groups (ANOVA main effect of virus p= 0.90; Fig. 4c). During the Test Phase, CNO-treated D1-hM4Di mice showed a trend level difference in choices to criterion (p=0.08 Mann-Whitney U test; Fig. 5c) and total nonrewarded choices compared to CNO-treated D1-mCherry mice (p=0.10 Mann-Whitney U test; Fig. 5d). D1-hM4Di mice were also slower to accumulate rewards to D1-mCherry mice (**p<0.01; Fig. 5e), and made significantly more choices to the training-naive preferred odor, O2, suggesting failure to implement learned suppression was affected by DMS dSPN inhibition (*p<0.05 Mann-Whitney U test; Fig. 5f). These data suggest that inhibiting DMS dSPNs can significantly impact Test Phase behavior and has effects similar to iSPN activation.

**Fig. 5.**
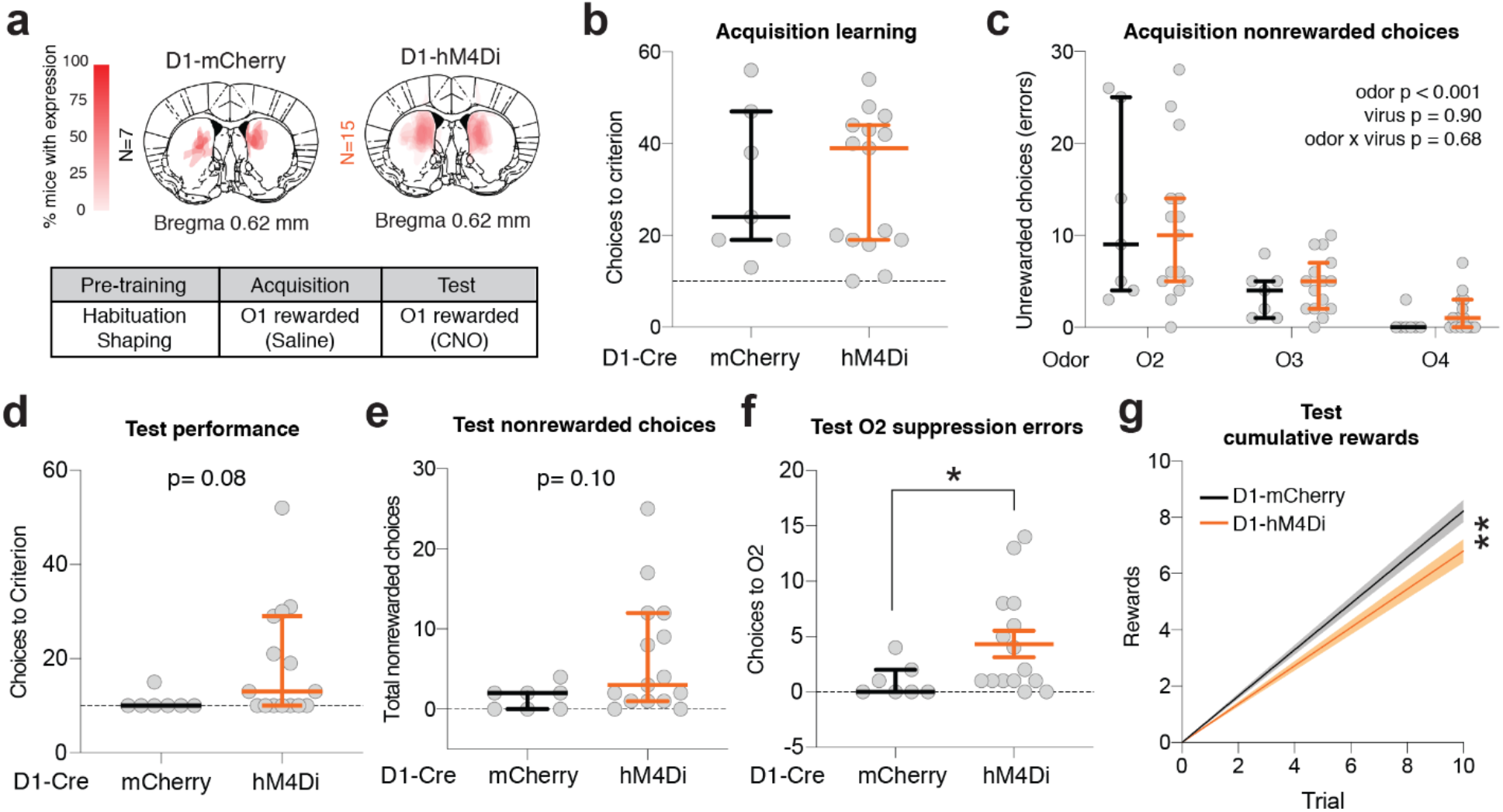
Chemogenetic manipulation data show dSPN inhibition impairs suppression of nonrewarded choices. **(a)** Top panel: schematic illustrating injection site and viral spread D1-Cre DIO-mCherry (N=7) and DIO-hM4Di (N=15) mice. Opacity indicates the number of mice in each cohort that expressed virus in a given location. Bottom panel: summary of behavior. **(b)** There was no significant difference between groups in Acquisition Phase choices to criterion (p=0.99 Mann-Whitney U test). **(c)** There was a significant effect of odor identity (F(1.1, 22.3) = 26.6, **p<0.0001; two-way ANOVA) but not virus (F(1,20)=0.02, p=0.90; two-way ANOVA) on the non rewarded choices made during Acquisition. **(d)** D1-hM4Di mice did not require significantly more choices to reach Test Phase criterion compared to D1-mCherry mice (p=0.08 Mann-Whitney test). **(e)** Total Test Phase nonrewarded choices did not significantly differ between D1-hM4Di and D1-mCherry groups (p=0.10 Mann-Whitney test). **(f)** D1-hM4Di mice failed to suppress choice of O2, the training-naive preferred odor, more often in the Test Phase compared to D1-mCherry mice (*p<0.05 Mann-Whitney test). **(g)** D1-hM4Di mice were significantly slower to accumulate rewards during Test Phase compared to D1-mCherry mice (**p<0.01). Mean ± SEM are plotted for normally distributed data including the D1-hM4Di group in panel g. Otherwise data plotted indicate Median ± IQR. Data in f indicate linear regression line with bands plotting the 95% confidence interval.

### Trial-by-trial RL modeling suggests that activating iSPNs alters Test Phase performance by increasing choice stochasticity

To further examine how chemogenetic manipulation affected underlying choice processes, we compared multiple reinforcement learning (RL) models (Daw, 2009) fit to trial-by-trial changes in behavior using a hierarchical fitting process (Fig. 6a; Methods). We did not include the biologically plausible OpAL model we used to generate predictions (Fig. 1, S1) in the candidate models fit to the data, because it is too complex to allow for satisfactory model fitting of the data in this task. Instead, we opted to fit and compare simpler RL models to both real mouse behavior and simulated OpAL behavior. We were thus able to validate the interpretation of the fit RL parameters of our best fit model by analyzing how it captures the behavior of simulated OpAL model. In our RL models, the process of learning the values of each odor (Q values) was adjusted by reward prediction errors and was separated from a selection process that transforms odor values into choice probabilities (see Methods for more details). The best fit model for our behavioral data included phase-specific parameters for the learning rate α and the inverse temperature parameter β which captures choice stochasticity (see Table 1 for alternate model comparison). In D2-Cre mice, we found that Acquisition Phase α and β parameters did not differ across groups (Fig. 6b), whereas the Test Phase β parameter was significantly lower for mice expressing activating DREADD in iSPNs (D2-hM3Dq) compared to mCherry control and inhibitory DREADD (D2-hM4Di) (Fig. 6c). These data suggest that chemogenetic activation of iSPNs in DMS makes choice policy more stochastic/exploratory. Model fits of OpAL-simulated trial histories converged on the same results, with simulated iSPN activation associated with decreased Test Phase β (Fig. S3). RL model fits to OpAL-simulated data also suggested that inhibition of dSPNs reduced the Test Phase β parameter, but RL fits to D1-hM4Di mice were not significantly different from D1-mCherry mice (Fig. 6d-f, S3). This may have been driven in part by the lower number of D1-mCherry mice.

**Fig. 6.**
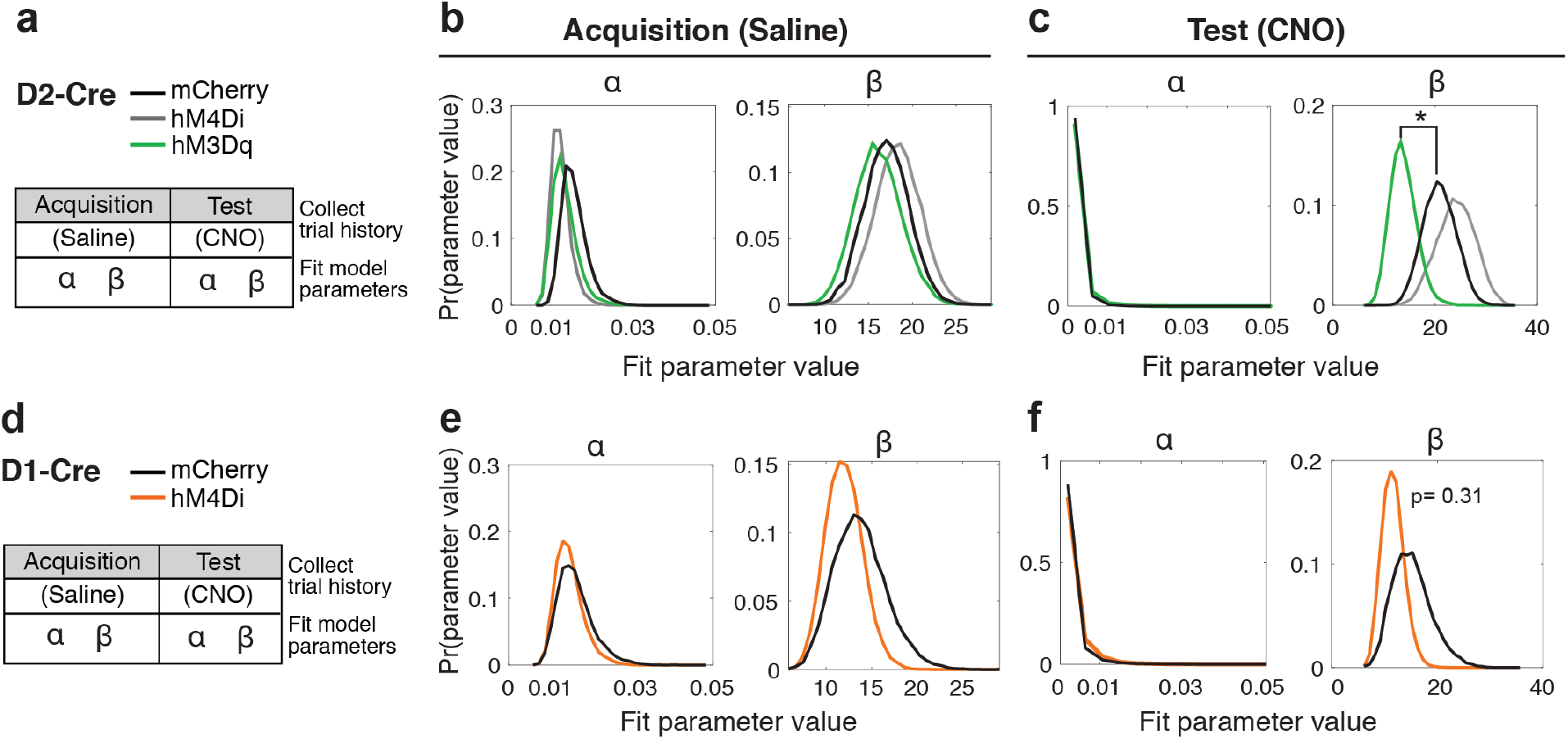
iSPN activation increases choice stochasticity. (**a**) Acquisition and Test Phase trial history data from D2-Cre DREADD mice and controls were modeled using an RL model, and best fit parameters were inferred using hierarchical Bayesian model fitting. (**b**) Test Phase alpha and β parameters did not significantly differ among manipulation groups (p>0.05). (**c**) Test Phase β was significantly lower in the D2-hM3Dq group compared to mCherry control (*p<0.05). (**d**) Acquisition and Test Phase trial history data from D1-Cre hM4Di mice and controls were modeled using an RL model, and best fit parameters were inferred using hierarchical Bayesian model fitting. **(e)** Test Phase alpha and β parameters did not significantly differ among manipulation groups (p>0.05 Kruskal Wallis ANOVA). **(f)** Test Phase β was not significantly different in the D1-hM4Di group compared to mCherry control (p=0.31). Hypothesis tests were conducted using Kruskal-Wallis ANOVA.

**Table 1.** RL model comparison.

| model | Learning rate $\alpha$ | Softmax $\beta$ | Phase decay | Number of parameters per participant | WAIC |
| --- | --- | --- | --- | --- | --- |
| ab | single | single | - | 2 | 5682 |
| abf | single | single | x | 3 | 5670 |
| abb | single | per phase | - | 3 | 5663 |
| aabf | per phase | single | x | 4 | 5635 |
| **aabb** | **per phase** | **per phase** | - | **4** | **5622** |
| a0bb | per phase; $\alpha_{\text{recall}}=0$ | per phase | - | 3 | 5916 |

We tested multiple instantiations of the model to capture task phase effects. Specifically, we investigated whether behavior would be better captured by shared or separate learning rate and softmax inverse temperature across phases; as well as whether decaying learned Q-values toward the initial Q-values between the two phases in proportion to a decay parameter could account for forgetting between the two phases (indicated by x). Last, we tested whether fixing learning rate to 0 in the recall phase would fit better. We used WAIC (Vehtari et al., 2017) for model comparison. We found that the best fitting model included a separate learning rate and softmax inverse temperature for both phases (aabb; displayed in bold text).

### Acute chemogenetic manipulation alters choice strategy in a manner not explained by locomotor effects

Because the DMS is associated with multiple aspects of behavior as well as decision-making, we also closely examined trial-by-trial behavior during Test Phase to evaluate potential effects of iSPN and dSPN chemogenetic manipulation on exploratory locomotion, impulsivity and motivation.

#### Quadrant “entries” as a metric of exploratory locomotion in the task arena

During each trial of the odor-based serial choice task, mice were free to enter each of the 4 quadrants and sample the odors present in each pot, quantified as entries, before making a bi-manual dig to indicate choice (Fig. 7a). Mice expressing activating DREADD in iSPNs (D2-hM3Dq) or inhibitory DREADD in dSPNs (D1-hM4Di) consistently made fewer entries during the Test Phase compared to mCherry controls (ANOVA main effect **p<0.01; Fig. 7b). Both D2-hM3Dq and D1-hM4Di mice were more likely to choose the first odor they encountered (classified as single entry trials) compared to mCherry control mice (ANOVA main effect ***p<0.0001; Fig. 7c). D2-hM3Dq and D1-hM4Di mice did not significantly differ from each other in Test Phase entries (Dunn’s multiple comparisons test p>0.99) or fraction single entry choices (Dunn’s multiple comparisons test p>0.99).

**Fig. 7.**
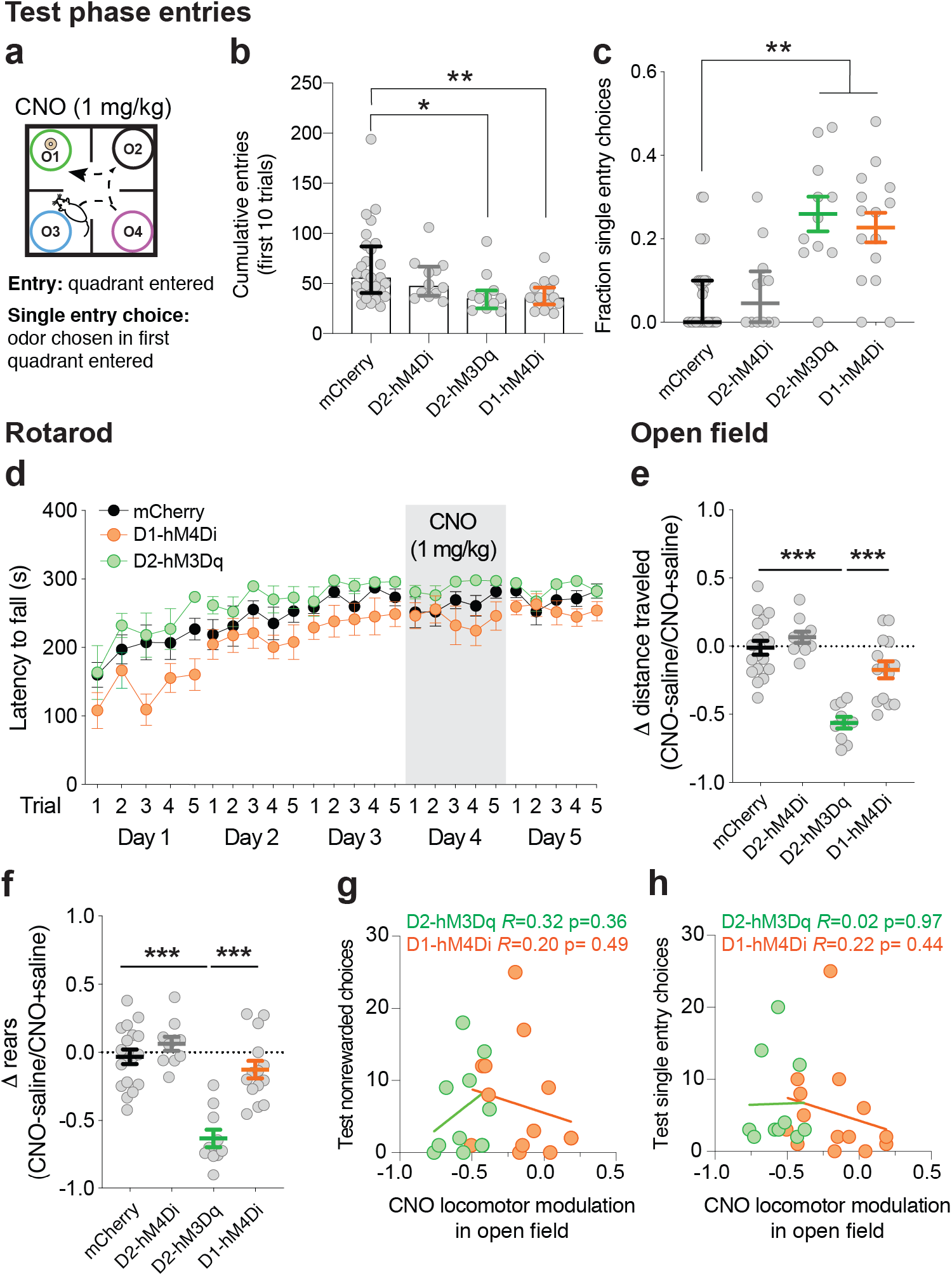
Chemogenetic manipulation of DMS direct and indirect pathway neurons reduced entries and increased single trial choices. Outside of the task context, chemogenetic manipulation spared rotarod performance but did affect open field locomotion in a manner uncorrelated with task choice effects. (**a**) Test Phase quadrant entries. (**b**) D2-Cre mice expressing activating DREADD (D2-hM3Dq) or D1-Cre mice expressing inhibitory DREADD (D1-hM4Di) made fewer entries during Test Phase on CNO compared to mCherry control mice on CNO (*p<0.05, **p<0.01 Kruskal Wallis ANOVA). (**c**) D2-hM3Dq and D1-hM4Di mice made significantly more single entry trials (F(3,40.65)= 9.55, ***p< 0.0001 main effect of group Brown-Forsythe ANOVA; **p<0.01 unpaired t-test with Welch’s correction). (**d**) Chemogenetic activation of iSPNs (D2-hM3Dq) or inhibition of dSPNs (D1-hM4Di) did not significantly affect latency to fall on the rotarod after stable performance was attained (main effect of drug compared across days 3-5 of stable performance F(1.98, 43.47) = 0.595, p=0.55, two-way repeated measures ANOVA). (**e**) Spontaneous locomotion was significantly reduced in D2-hM3Dq mice on CNO (1 mg/kg) compared to saline (F(3,48)= 22.38, ***p<0.0001, one-way ANOVA, ***p<0.0001, uncorrected Fisher’s LSD) and D1-hM4Di mice to a lesser extent (*p<0.05, uncorrected Fisher’s LSD). (**f**) CNO administration significantly reduced the number of vertical rears made by D2-hM3Dq mice (F(3,48)= 21.87 ***p<0.0001, one-way ANOVA, ***p<0.0001 mCherry vs. D2-hM3Dq uncorrected Fisher’s LSD). (**g**) Locomotor modulation on CNO (presented in e) did not correlate with the number of choices of nonrewarded odors during Test Phase for either D2-hM3Dq or D1-hM4Di groups. (**h**) Locomotor modulation on CNO (presented in e) did not correlate with the number of single entry trials mice made on CNO during Test Phase for either D2-hM3Dq or D1-hM4Di groups.

#### Random or value-ranked choice as a metric of impulsiveness

There was a significant effect of odor identity on single entry choices that tracked with the odor values learned during Acquisition (ANOVA main effect ***p<0.0001; Fig. S4), suggesting that single entry choices were guided by learned odor values and were not simply impulsive. The rewarded odor, O1, and the training-naive preferred O2 were the two most frequently chosen odors for single entry trials (Fig. S4).

#### Trial omissions and total trials as metrics of motivation

D2-hM3Dq and D1-hM4Di mice did not make more omissions (3 minute trial without making a choice) compared to mCherry controls (ANOVA main effect p=0.08; Fig. S4), and they completed the task to criterion (completing the same number or more trials than controls) (Fig. 4d; Fig. 5d), suggesting that they remained highly motivated in the task.

#### Locomotion outside of the task

To examine the effects of DMS DREADD manipulations on locomotor behavior more generally, we measured the effect of CNO on rotarod performance and spontaneous locomotion ~2 weeks after mice completed the odor-guided serial choice task. CNO did not alter latency to fall in the rotarod in any groups tested (Fig. 7d). CNO administration did reduce spontaneous locomotion in the open field in mice expressing activating DREADD in iSPNs (D2-hM3Dq) compared to mCherry control mice, measured as change in distance traveled (Tamhane’s T2 multiple comparisons test ***p<0.0001) and change in the number of vertical rears (Tamhane’s T2 multiple comparisons test ***p<0.0001) on CNO vs. saline. Meanwhile, spontaneous locomotion of mice expressing inhibitory DREADD in iSPNs (D2-hM4Di) or dSPNs (D1-hM4Di) did not significantly differ from mCherry control mice on CNO (D2-hM4Di vs. mCherry p= 0.69; D1-hM4Di vs. mCherry p= 0.20; Fig. 7e,f). It is worth noting that while the suppressing effect of CNO on spontaneous locomotion was greater in D2-hM3Dq mice compared to D1-hM4Di mice (Tamhane’s T2 multiple comparisons test ***p<0.001), both groups exhibited a similar reduction in total entries (Dunn’s multiple comparisons test p>0.99) and increase in single entry trials (Dunn’s multiple comparisons test p>0.99) within the serial choice task (Fig. 7b,c). The magnitude of CNO’s effect on spontaneous locomotion did not correlate with nonrewarded choices during Test or single entry trials (Fig. 7h). Finally, single entry trial latencies did not differ across groups, suggesting that they were qualitatively similar when made by D2-hM3Dq or D1-hM4Di mice and controls (Fig. S4). Together these data suggest that the influence of CNO on choice behavior in the Test Phase was not simply motor-related.

## Discussion

In the present study, we found that manipulating iSPN activity had effects on learned choice behavior that were contrary to a prominent ‘select/suppress’ heuristic model. We showed that chemogenetically activating iSPNs or inhibiting dSPNs impaired suppression of nonrewarded choices, whereas inhibiting iSPNs did not affect choice behavior. These behavioral results were predicted by the OpAL network model, in which learned values are adjusted in the direct and indirect pathway by an update rule that emphasizes positive and negative prediction errors in direct and indirect pathways, respectively. Choice values are then compared after the weighted difference of direct and indirect pathway activity has been calculated (Collins and Frank, 2014). RL model fits to OpAL simulated data and mouse behavioral data showed that activating iSPNs reduced the inverse temperature (β) parameter, consistent with more exploratory choice. In the context of our deterministic task, this manifested as failure to suppress nonrewarded choices. Our computational and empirical data demonstrate that the combined output of direct and indirect pathways, and not the independent function of either, is critical for learned choice suppression.

Previous studies provide clear evidence that optogenetic stimulation of DMS iSPNs can drive aversion and inhibit movement (Kravitz et al., 2010; Kravitz et al., 2012; Freeze et al., 2013; Roseberry et al., 2016), suggesting that choice suppression might be an extension of indirect pathway ‘no go’ function. However, recording data collected during decision-making has shown that dSPNs and iSPNs are simultaneously active when animals choose an action (Cui et al., 2013; Isomura et al., 2013; Jin et al., 2014). The ‘select/suppress’ heuristic accounts for this coactivation by suggesting that dSPNs select specific actions while iSPNs simultaneously suppress alternate actions. The OpAL model can also accommodate coactivation but suggests that the choice is a function of the weighted difference of activity in the two pathways and not the independent action of either pathway.

While many studies have focused on the role of the striatum in selecting rewarded actions (Lau and Glimcher, 2008; Tai et al., 2012; Nonomura et al., 2018) and stimuli (Santacruz et al., 2017), fewer have studied its role in avoiding low-value actions (Bryden et al., 2012; Schmidt et al., 2013; Friedman et al., 2015; Ogasawara et al., 2018) and stimuli (Kim et al., 2017). Here, our goal was to understand how activity in DMS iSPNs and also dSPNs influences the ability to suppress an initially encountered low-value choice in order to make a subsequent high-value choice. In our odor-guided serial choice task, as in many natural decision-making settings, the value of a given action/choice (here, dig) was contingent on available stimulus information plus choice and outcome history. RL model fits to odor choice trial histories enabled us to investigate how chemogenetic manipulation altered the relationship between odor value estimates and choice. The multiple choice task design enhanced our ability to interpret underlying choice processes. The fact that the mice were freely moving and were never required to hold still removed a common confound between choice suppression and motor freezing. Lastly, the task was acquired in a single session without extensive training that is often found in rodent operant tasks. This should enhance relevance to DMS function, which is engaged in early flexible goal-directed learning (Yin et al., 2009; Thorn et al., 2010; Kupferschmidt et al., 2017; Matamales et al., 2020). Collectively, these more ethological task features may have permitted novel observations about the role of the indirect pathway in choice suppression behavior (Juavinett et al., 2018).

In addition to updating our understanding of choice suppression, our data add new circuit dimension to previously proposed dopamine and D2R mechanisms underlying choice exploration. We found that when iSPNs were activated, choice became more stochastic/exploratory, meaning that mice were more likely to “explore” (i.e. choose) a lower value odor as opposed to “exploit” the highest value odor, as estimated by RL model fits. This was captured by a lower inverse temperature parameter, which tunes explore/exploit balance in the estimated odor value to choice conversion. This observation is consistent with a previous study that found D2R antagonism in the primate caudate reduced the inverse temperature parameter and increased exploratory choice (Lee et al., 2015). Our findings are also compatible with computational accounts that predict that lowering tonic dopamine, which facilitates iSPN activity and suppresses dSPN activity (Mallet et al., 2006; Surmeier et al., 2007), shifts explore/exploit balance towards exploration (Humphries et al., 2012; Dunovan and Verstynen, 2016). Finally, if behavioral switching is viewed as exploring action space, our iSPN data may relate to recent studies that report iSPN activity increases in response to outcomes preceding switch trials (Geddes et al., 2018; Nonomura et al., 2018).

Greater choice exploration did not go hand in hand with greater locomotor exploration. Unexpectedly, we observed that iSPN activation and dSPN inhibition increased the number of trials in which mice chose the first odor they encountered (Fig. 7). Single entry trial choices were significantly influenced by odor identity and odor Q values captured by the RL model (Fig. S4), suggesting that they were not impulsive or random. Chemogenetic manipulation did not affect rotarod performance and the effects of manipulation on spontaneous locomotion in the open field did not correlate with performance in the serial choice task (Fig. 7). Therefore, we interpret that these single entry trials are the result of choices made when the chemogenetic manipulation minimized the difference in choice weights across odors. This could be related to recent work in the basal ganglia on speed/accuracy tradeoffs (Mazzoni et al., 2007; Baraduc et al., 2013) and net expected return and state value signals (Wang et al., 2013; Hamid et al., 2016). Future experiments which manipulate alternative choice value and choice discriminability should further inform these observations.

Overall, our data support existing models of basal ganglia function in which trial and error choice drives learning that is later stored or read out in the activity emerging from DMS dSPNs and iSPNs (Bariselli et al., 2018), but inverts the often ‘expected’ direction of activity assumed to be important in the indirect pathway iSPNs for choice suppression. Notably, despite what some might consider a surprising outcome, our data are consistent with previous observations of synaptic plasticity following goal-directed action learning (Shan et al., 2014), which reveal potentiation onto dSPNs and depression onto iSPNs after learning. Further work will need to be done to inform how long term potentiation and depression are allocated to specific neural ensembles of dSPNs and iSPNs to sculpt choice.

In summary, our findings suggest that the indirect pathway does not independently mediate choice suppression. Instead, our model and data suggest choice arises from the weighted difference in dSPN and iSPN population activity, and conditions that reduce this difference increase choice stochasticity/exploration. Importantly, we demonstrate that manipulations that simply enhance activity in the indirect pathway do not facilitate adaptive choice suppression, and in fact can have the opposite effect. These data highlight the importance of using network concepts and models over simple heuristic accounts of circuit function to understand decision-making. We are hopeful that these findings will inform studies of addiction and other conditions in which greater capacity for choice suppression is desirable.

---

**Supplementary 1 associated with Fig. 1.**
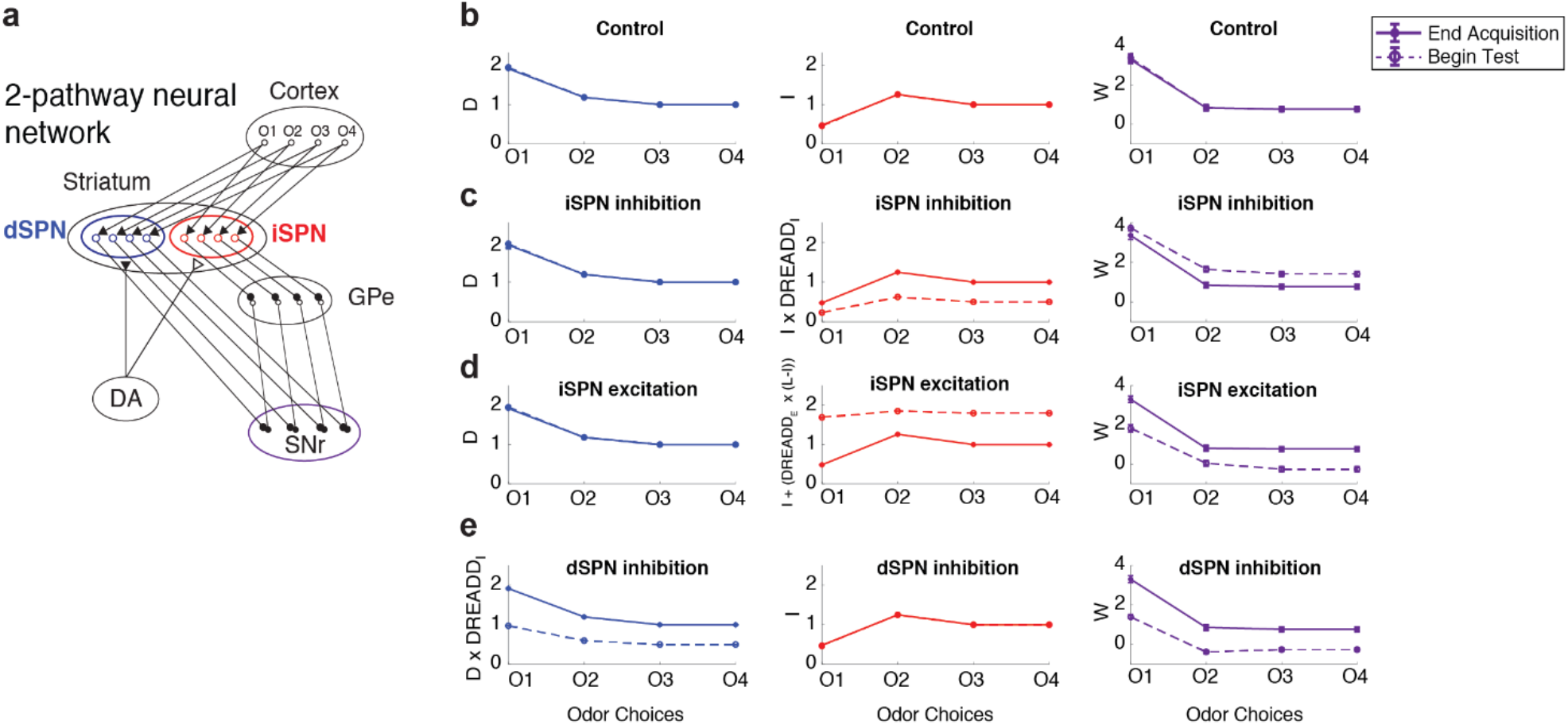
OpAL model simulations under control and iSPN, dSPN manipulation conditions. **(a)** Schematic of 2-pathway cortico-basal ganglia network. The 4 odor options are represented as separate action channels in the direct and indirect pathways. **(b)** Mimicking chemogenetic manipulation in OpAL model simulation. D and I weights for Odors (1-4) at the end of Acquisition learning are depicted in solid blue and red lines. D and I weights at the beginning of Test Phase (dashed line) do not differ in the unmanipulated control group. Choice weights (W), depicted in purple, are calculated as the weighted difference of D and I, which are each scaled by separate *β* parameters (*β*_*D*_ and *β*_*I*_). *β*_*D*_ has a larger dynamic range than *β*_*I*_, and consequently D weights have a greater influence on choice weights (W). **(c)** iSPN inhibition is implemented by multiplicatively reducing I weights (dashed red line) which alters choice weights (dashed purple line). **(d)** iSPN activation is implemented as an additive effect that is scaled by I minus an upper limit, L, that causes low I weights to increase more than high I weights (dashed red line) which alters choice weights (dashed purple line). **(e)** dSPN inhibition is implemented by multiplicatively reducing I weights (dashed blue line) which alters choice weights (dashed purple line).

**Supplementary 2 associated with Fig. 4.**
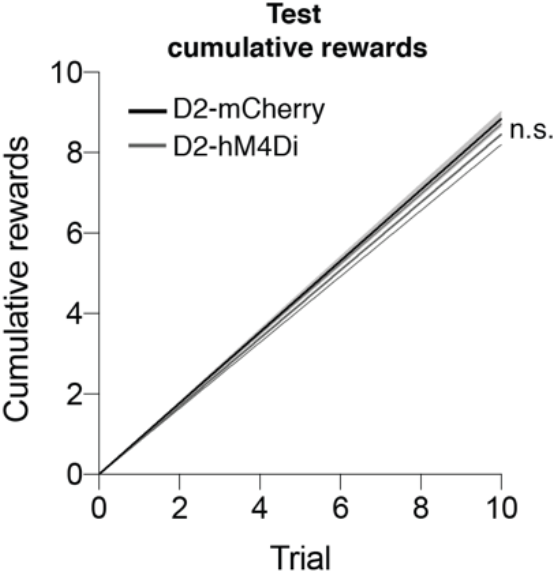
iSPN chemogenetic inhibition does not significantly affect Test Phase reward accumulation compared to D2-mCherry control mice. Linear regression lines fit to first 10 choices of Test Phase with 95% confidence bands plotted. D2-mCherry and D2-hM4Di regression slope estimates overlapped at the 95% confidence interval.

**Supplementary 3 associated with Fig. 6.**
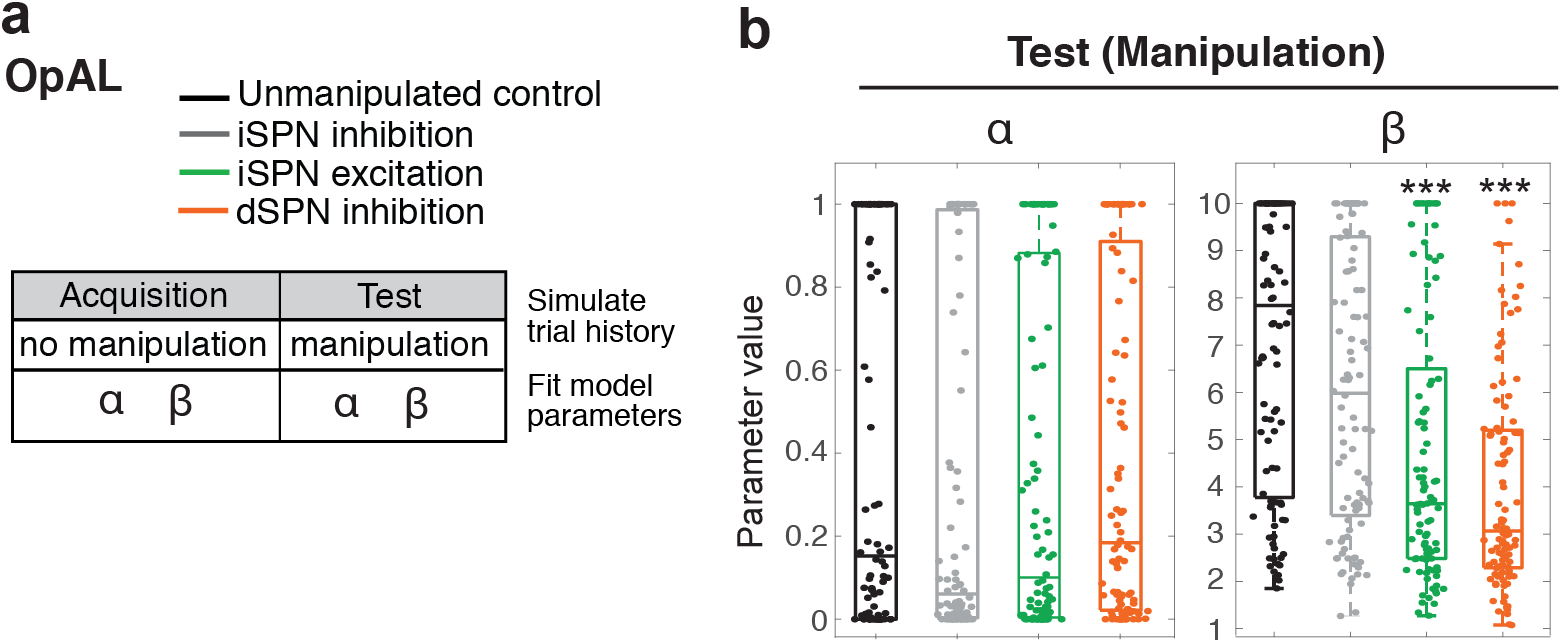
RL model fit of OpAL simulation data. **(a)** Trial histories were simulated using OpAL model with or without DREADD manipulation implemented during Test Phase. Simulated data was modeled using the same RL model as in Fig. 7 and best fit parameters were inferred using a standard, non-hierarchical fitting procedure. **(b)** Similar to the behavioral manipulation data, simulated iSPN activation data was captured by significantly smaller Test Phase beta compared to the unmanipulated group (***p<0.001). In addition, dSPN inhibition data was captured by a significantly smaller Test Phase beta compared to the unmanipulated group (***p<0.001). Hypothesis tests were conducted using Wilcoxon signed rank tests.

**Supplementary 4 associated with Fig. 7.**
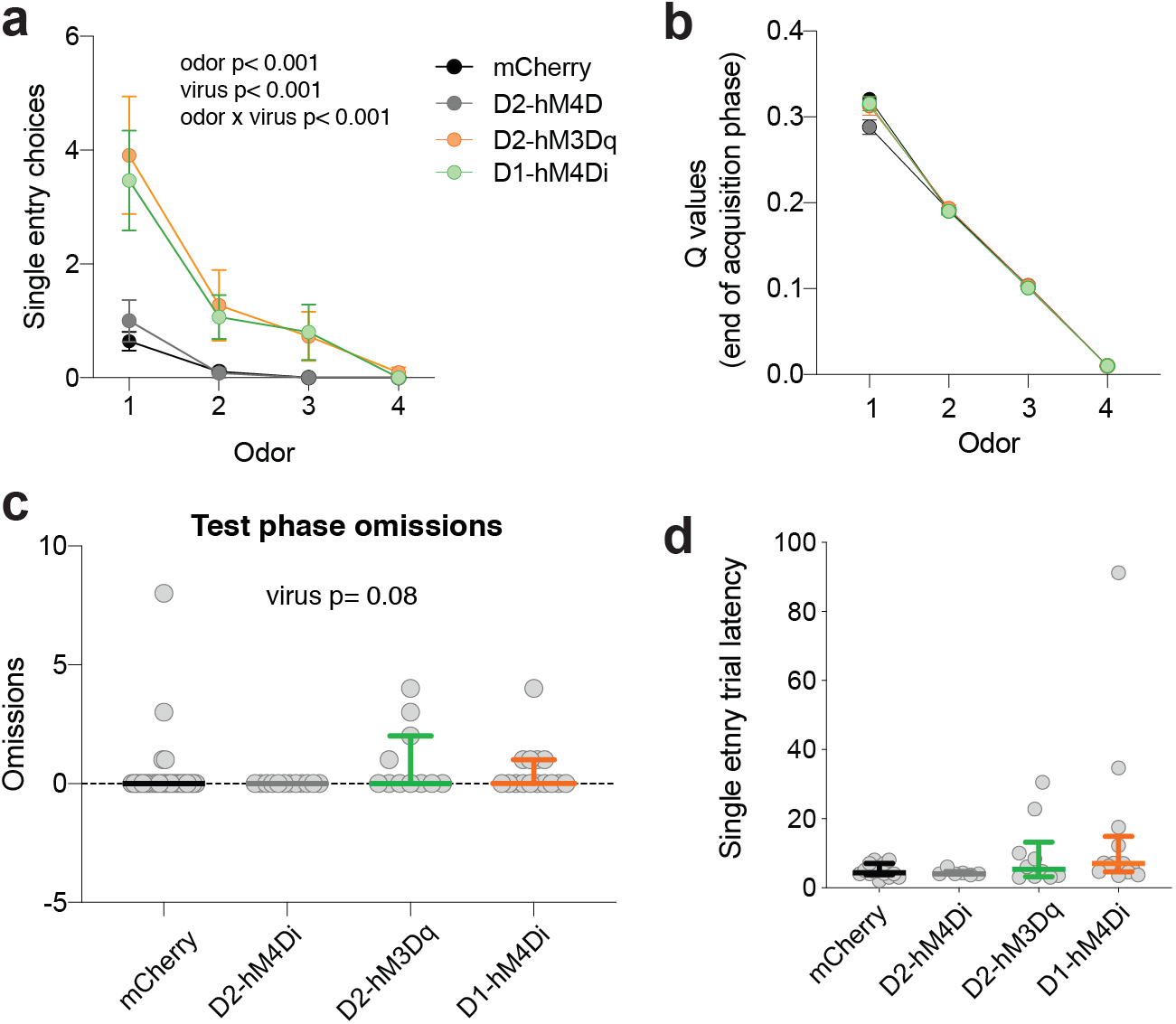
Single entry trials were related to odor value and were not explained by differences in motivation to work in the task. **(a)** There was a significant main effect of odor identity (F(1.58, 98.22)= 50.02 ***p<0.0001), virus (F(3, 62)= 7.3 ***p<0.001), and interaction (F(9, 186)= 7.1 ***p<0.0001) on the number of single entry choices during the Test Phase. **(b)** Odor Q values at the end of Acquisition for each group. **(c)** There was no significant effect of virus on Test Phase omissions (no choice made during 3 minute trial) (p=0.08; Kruskal-Wallis ANOVA). **(d)** There was no significant effect of virus on single entry trial latency (p=0.15; Kruskal-Wallis ANOVA). Data are presented as Mean ± SEM in panels a,b and Median ± IQR in panels c,d.

## Acknowledgments

We thank Yuting Zhang, Satya Vedula, Christopher Hall, and Nana Okada for assistance with behavior and histology. We thank Dr. Richard Ivry for manuscript feedback and Wilbrecht and Collins lab members for helpful discussion. This research was supported by a National Institute of Mental Health postdoctoral fellowship under Grant F32MH110184 to K.D.

## Author contributions

K.D., A.G.E.C., and L.W. designed the research. K.D. performed all experiments and analyzed data. A.G.E.C. contributed analytic tools, designed and performed OpAL simulation and RL model analysis to which B.H. contributed. K.D., A.G.E.C., and L.W. wrote the manuscript.

## Declaration of interests

The authors declare that there are not conflicts of interest.

## Methods

### OpAL model

OpAL is an actor-critic-like RL algorithm which assumes two sets of weights are being tracked, D and I (corresponding to the direct pathway and indirect pathways, respectively; initialized at 1.5 for the preferred odor D weight, 1 for all other weights), in addition to a classic critic value V (initialized at 0.25 as the initial expected value of each option). Critic values are updated according to the classic RL equation:

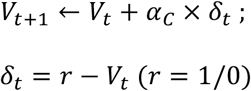

D and I weights are updated with a three factor, non-linear learning rule that emphasizes gains and losses, respectively:

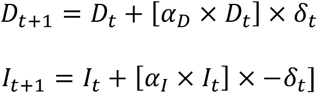

To simplify the modeling of excitatory DREADD effects, we enforce an upper limit L to the D and I weights.

The final choice is a softmax probability based on combined weights:

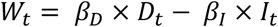

which is supposed to represent the output of the cortico-basal ganglia loops.

Chemogenetic tools modulate activity of dSPNs and iSPNs neurons, and we model their effects on D and I weights respectively. We assume that inhibitory DREADD multiplicatively decrease the corresponding pathway’s activity:

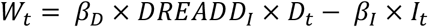

in the case of dSPN DREADD.

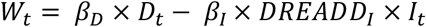

in the case of iSPN DREADD.

For the simulations, the inhibitory *DREADD*_*I*_ parameter is fixed at 0.5.

For excitatory DREADD, we assume that the tool promotes firing in neurons whose activity would otherwise be subthreshold, thus increasing low weights more than high weights. Specifically, we model *W* activity as:

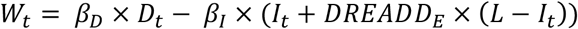

in the case of iSPN DREADD, where *L* is the activity limit. For all simulations, *L* is fixed at 2, and the excitatory *DREADD*_*E*_ parameter is fixed at 0.8.

To investigate the effects of chemogenetic manipulation on behavior, we simulated 100 times with parameters set to *α*_*C*_ = *α*_*D*_ = *α*_*I*_ = 0.1, randomly chosen parameters 1 < *β*_*D*_ < 3, and 1 < *β*_*I*_ < 1.6, reflecting greater influence of the direct than indirect pathway on the final choice.

### Mice

All mice were weaned on postnatal day (P)21 and group-housed on a 12:12hr reverse light:dark cycle (lights on at 10PM). C57BL/6 BAC transgenic mice expressing Cre recombinase under the regulatory elements for the D1 and D2 receptor (Drd1a-Cre and D2-Cre ER43) were obtained from Mutant Mouse Regional Resource and bred in our colony. Mice had ad lib access to food and water before food restriction in preparation for training. All procedures were approved by the Animal Care and Use Committee of the University of California, Berkeley and complied with the NIH guide for the use and care for laboratory animals.

### Viruses and tracers

Adeno-associated viruses (AAVs) were produced by the Gene Therapy Center Vector Core at the University of North Carolina at Chapel Hill or by Addgene viral service and had titers of >10^12^ genome copies per mL. For chemogenetic manipulations, mice were bilaterally injected with 0.5 uL of rAAV8-hsyn-DIO-mCherry, rAAV8-hsyn-DIO-hM3Dq-mCherry, or rAAV8-hsyn-DIO-hM4Di-mCherry. For *in vitro* electrophysiological validation experiments of rAAV8-hsyn-DIO-hM4Di-mCherry, mice were bilaterally injected with 0.69 uL of a 2:1 mixture of rAAV8-hsyn-DIO-hM4Di-mCherry and rAAV5-Ef1α-DIO-hChR2-EYFP.

### Stereotaxic Virus Injection

Male and female mice (6-8 weeks) were deeply anesthetized with 5% isoflurane (vol/vol) in oxygen and placed into a stereotactic frame (Kopf Instruments; Tujunga, CA) upon a heating pad. Anesthesia was maintained at 1-2% isoflurane during surgery. An incision was made along the midline of the scalp and small burr holes were drilled over each injection site. Virus or tracer was delivered via microinjection using a Nanoject II injector (Drummond Scientific Company; Broomall, PA). Injection coordinates for DMS were (in mm from bregma): 0.90 anterior, +/−1.4 lateral, and −3.0 from surface of the brain. Injection coordinates for SNr were: 3.2 posterior, 1.2 lateral, and −4.6 from the surface of the brain. Mice were given subcutaneous injections of meloxicam (10 mg/kg) during surgery and 24 & 48 hours after surgery. Mice were group-housed before and after surgery and 4-6 weeks were allowed for viral expression before behavioral training or electrophysiology experiments.

### Drugs

Clozapine-N-Oxide was generously provided by the NIMH Chemical Synthesis and Drug Supply Program (NIMH C-929). CNO was made fresh each day and dissolved in DMSO (0.5% final concentration) and diluted to 0.1 mg/mL in 0.9% saline USP. Tetrodotoxin (TTX), D-AP5, and NBQX disodium salt were purchased from Tocris Biosciences (Ellisville, MO).

### Electrophysiology

Mice were deeply anesthetized with an overdose of ketamine/xylazine solution and perfused transcardially with ice-cold cutting solution containing (in mM): 110 choline-Cl, 2.5 KCl, 7 MgCl2, 0.5 CaCl2, 25 NaHCO3, 11.6 Na-ascorbate, 3 Na-pyruvate, 1.25 NaH2PO4, and 25 D-glucose, and bubbled in 95% O2/ 5%CO2. 300 μm thick sections (sagittal for optogenetic stimulation experiment, coronal for all others) were cut in ice-cold cutting solution before being transferred to ACSF containing (in mM): 120 NaCl, 2.5 KCl, 1.3 MgCl2, 2.5 CaCl2, 26.2 NaHCO3, 1 NaH2PO4 and 11 Glucose. Slices were bubbled with 95% O2/ 5% CO2 in a 37°C bath for 30 min, and allowed to recover for 30 min at room temperature before recording. All recordings were made using a Multiclamp 700B amplifier and were not corrected for liquid junction potential. The bath was heated to 32°C for all recordings. Data were digitized at 20 kHz and filtered at 1 or 3 kHz using a Digidata 1440 A system with pClamp 10.2 software (Molecular Devices, Sunnyvale, CA, USA). Only cells with access resistance of <25 MΩ were retained for analysis. Access resistance was not corrected. Cells were discarded if parameters changed more than 20%. Data were analyzed using pClamp or R (RStudio 0.99.879; R Foundation for Statistical Computing, Vienna, AT).

Spontaneous spiking in GPe neurons was recorded in cell-attached configuration. To evoke synaptic transmission by activating ChR2, we used a single wavelength LED system (470 nm; Thorlabs; Newtown, NJ) connected to the epifluorescence port of the Olympus BX51 microscope. Light pulses of 1-10 ms triggered by a TTL (transistor-transistor logic) signal from the Clampex software (Molecular Devices; Sunnyvale, CA) were delivered through a 63x objective and used to evoke synaptic transmission. Blue light pulses were delivered once every 10 s, and a minimum of 30 trials were collected. Light-evoked IPSCs were recorded in whole-cell configuration at +10 mV holding potential in the presence of D-AP5 (50 μM) and NBQX disodium salt (33 μM) to block glutamatergic neurotransmission. Recording pipettes had 2.5-5.5 MΩ resistances and were filled with internal solution (in mM): 115 Cs-methanesulfonate, 10 HEPES, 10 BAPTA, 10 Na2-phosphocreatine, 5 NaCl, 2 MgCl2, 4 Na-ATP, 0.3 Na-GTP.

Whole-cell current clamp recordings were performed using a potassium gluconate-based intracellular solution (in mM): 140 K Gluconate, 5 KCl, 10 HEPES, 0.2 EGTA, 2 MgCl2, 4 MgATP, 0.3 Na2GTP, and 10 Na2-Phosphocreatine. For current clamp recordings to validate CNO induced depolarization in Gq-DREADD-expressing Drd2+ neurons, ACSF contained 0.5 μM TTX and a stable baseline was collected for 3-5 minutes before ACSF containing 0.5 μM TTX + 10 μM CNO was washed on. For all electrophysiology experiments, both male and female mice were used.

### Behavioral assays

Adult male and female mice (6-10 weeks) were used in behavioral assays. Mice were first tested in 4 choice odor-guided serial choice task and then ≥ 2 weeks later were tested in locomotor and/or rotarod tasks so that performance on CNO could be compared within animals across tasks. Prior to all behavior assays, mice were habituated to the testing room for 30 minutes, and all behavior testing began 30 min after CNO treatment. Importantly, all groups (including DIO-mCherry) were administered CNO to control for potential off-target effect of the CNO metabolite clozapine (Mahler and Aston-Jones, 2018).

### 4 choice odor-guided serial choice task

The odor-guided serial choice task used has previously been described in detail (Johnson and Wilbrecht, 2011; Johnson et al., 2016). In this task only the odor cue is predictive, and spatial or egocentric information are irrelevant. This behavior is also ethologically relevant because mice use odor information to locate food sources (Howard et al., 1968). Briefly, mice were food restricted to ~85 % bodyweight prior to training. On day 1, mice were habituated to the testing arena, on day 2 were taught to dig for cheerio reward in a pot filled with unscented wood shavings, on day 3 underwent a 4-choice odor discrimination in which they acquire the rule that 1 of 4 presented odors is rewarded, and finally on day 4 were tested for recall of the previously learned odor-reward association (Fig. 2a). During the Test Phase of the task, mice learned to discriminate among four pots with different scented wood shavings (anise, clove, litsea and thyme). All 4 pots were sham-baited with cheerio (under wire mesh at bottom) but only one pot was rewarded (anise). The pots of scented shavings were placed in each corner of an acrylic arena (12”, 12”, 9”) divided into 4 quadrants. Mice were placed in a cylinder in the center of the arena, and a trial started when the cylinder was lifted. Mice were then free to explore the arena until a choice was signaled by a dig to the wood shavings. The cylinder was lowered as soon as a choice was made. If the choice was incorrect, the trial was terminated and the mouse was gently encouraged back into the start cylinder. Trials in which no choice was made within 3 minutes were considered omissions. If mice omitted for 2 consecutive trials, they received a reminder: a baited pot of unscented wood shavings was placed in the center cylinder and mice dug for the “free” reward. Mice were disqualified if they committed 4 pairs of omissions. The location of the 4 odors was shuffled on each trial, and criterion was met when the mouse completed 8 out of 10 consecutive trials correctly. 24-hours after completing the Acquisition Phase, mice underwent a recall Test of the initial odor-reward rule to criterion. For chemogenetic manipulation experiments, mice were injected with saline 30 minutes prior to Acquisition training and injected with CNO (1.0 mg/kg) 30 minutes prior to the Test Phase. During Acquisition and Test Phase, experimenters (blind to group) manually scored entries into each quadrant, latency to dig, and odor choices. mCherry mice (D2-mCherry or D1-mCherry) were run in parallel with DREADD mice of the same genotype.

### Rotarod test

Females and males were run during separate sessions. On day 1, mice underwent a habituation trial in which they were placed individually in a clean holding cage for 5 mins. The rotarod (47650 Rota-Rod NG Ugo Basile; Monvalle VA, Italy) was then set at 5 rpm constant speed and each mouse was placed on the rod for 1 minute. The mice were then returned to the holding cage for another 5 mins before initiating the first trial. Each session consisted of 5 trials in which the rotarod constantly accelerated from 5-40 rpm over a period of 300 secs, and the latency at which mice fell off the or held onto the rod for a full rotation was recorded. Mice rested for 5 mins in the holding cage between trials. Asymptotic performance was reached by day 3 of training (Fig. 7). On day 4, DIO-DREADD and DIO-mCherry mice were administered CNO (1 mg/kg, i.p.) 30 minutes before rotarod testing began. On day 5 mice were tested drug-free in rotarod performance. The rotarod apparatus was cleaned between mouse cohorts with 3% hydrogen peroxide (for plastic components) and 70% ethanol (for metal troughs).

### Open field locomotor assay

On day 1, mice underwent a habituation session in which they were placed in a clear acrylic box (225 × 225 mm) inside a sound attenuated chamber (Med Associates; Fairfax, VT) with lights off. Locomotion was monitored for 15 minutes using infrared beam breaks (Versamax, AccuScan Instruments, Columbus, OH). On days 2 and 3 mice received injections of saline or CNO (counterbalanced across mice) 30 minutes before their locomotion was monitored for 15 minutes. The chamber was cleaned with 70% ethanol between mice.

### Histology

Mice were transcardially perfused with PBS followed by 4% PFA in PBS. Following 24h postfixation, coronal brain slices (75 μm) were sectioned using a vibratome (VT100S Leica Biosystems; Buffalo Grove, IL). To confirm viral targeting, we performed a standard immunohistochemical procedure using a primary antibody against red fluorescence protein (RFP) (rabbit, Rockland 600-401-379; 1:1000) to enhance the mCherry signal expressed in mice transduced with rAAV8-hSyn-DIO-DREADD-mCherry or rAAV8-hSyn-DIO-mCherry. Sections were counterstained with DAPI (Life Technologies; Carlsbad, CA). Images were acquired with a Zeiss Axio Scan.Z1 epifluorescence microscope (Molecular Imaging Center, UC Berkeley) at 10x magnification and viewed using FIJI (ImageJ). For colocalization experiments, mCherry signal was enhanced as previously described, and images were acquired using a Zeiss LSM 710 confocal microscope (Biological Imaging Facility, UC Berkeley). Anatomical regions were identified according to the Mouse Brain in Stereotaxic Coordinates by Franklin and Paxinos and the Allen Institute Mouse Brain Atlas.

### RL model

We modeled Acquisition and Test Phase behavior using a reinforcement learning model driven by an iterative error-based rule (Rescorla and Wagner, 1972; Sutton and Barto, 1998). The model uses a prediction error (δ) to update the value (V) of each odor stimulus, where δ is the difference between the experienced feedback (λ) and the current expected value (λ= 100 for rewarded, λ= 0 for unrewarded) scaled by a learning rate parameter (α), with 0<α<1. Because mice exhibit innate preferences for odors, we set initial odor values to fixed parameters [v1,v2,v3,v4] for all 75 mice tested.

To model trial-by-trial choice probabilities, the stimulus values were transformed using a softmax function to compute the relative probability of each choice. The inverse temperature parameter (β) determined the stochasticity of the choices. We used hierarchical Bayesian model fitting to infer the best fitting parameters, using the package STAN in Matlab (Carpenter et al., 2017). We assumed that odor values were shared by all animals, and that other parameters (α and β for each phase) were drawn from group level distributions defined by the experimental manipulation. We performed statistical tests on the distribution of samples obtained for the group-level hyperparameters. We compared the alternative models using the WAIC (Watanabe, 2010; Vehtari et al., 2017) and found that the best fit model included phase-specific (non-zero) α and β parameters; all RL model comparisons are presented in Table S1.

### Statistics

Statistical tests were performed in GraphPad Prism 7.0 (San Diego, CA) and the R programming environment. Groups were compared using one-way ANOVA if data were normally distributed or Kruskal-Wallis ANOVA if data were not normally distributed. When the ANOVA yielded significant results (p<0.05), a post-hoc Tamhane’s T2 test or Dunn’s test was used to compare DREADD manipulation groups to the mCherry control group. In several cases, D2-hM3Dq and D1-hM4Di groups were also compared (Fig. 7). All hypothesis testing was corrected for multiple comparisons. Data from D2-mCherry and D1-mCherry groups were pooled into a single ‘mCherry’ group for locomotor analysis presented in Fig. 7.

## Notes

### Competing Interest Statement

The authors have declared no competing interest.

## References

Albin RL, Young AB, Penney JB (1989) The functional anatomy of basal ganglia disorders. Trends in neurosciences 12:366–375.

Alexander GE, Crutcher MD (1990) Functional Architecture of Basal Ganglia Circuits – Neural Substrates of Parallel Processing. Trends in neurosciences 13:266–271.

Balleine BW, O’Doherty JP (2010) Human and rodent homologies in action control: corticostriatal determinants of goal-directed and habitual action. Neuropsychopharmacology : official publication of the American College of Neuropsychopharmacology 35:48–69.

Balleine BW, Delgado MR, Hikosaka O (2007) The role of the dorsal striatum in reward and decision-making. The Journal of neuroscience : the official journal of the Society for Neuroscience 27:8161–8165.

Baraduc P, Thobois S, Gan J, Broussolle E, Desmurget M (2013) A Common Optimization Principle for Motor Execution in Healthy Subjects and Parkinsonian Patients. Journal of Neuroscience 33:665–677.

Barbera G, Liang B, Zhang L, Gerfen CR, Culurciello E, Chen R, Li Y, Lin DT (2016) Spatially Compact Neural Clusters in the Dorsal Striatum Encode Locomotion Relevant Information. Neuron 92:202–213.

Bariselli S, Fobbs WC, Creed MC, Kravitz AV (2018) A competitive model for striatal action selection. Brain Res.

Bryden DW, Burton AC, Kashtelyan V, Barnett BR, Roesch MR (2012) Response inhibition signals and miscoding of direction in dorsomedial striatum. Front Integr Neurosci 6:69.

Carpenter B, Gelman A, Hoffman MD, Lee D, Goodrich B, Betancourt M, Riddell A, Guo JQ, Li P, Riddell A (2017) Stan: A Probabilistic Programming Language. J Stat Softw 76:1–29.

Castane A, Theobald DE, Robbins TW (2010) Selective lesions of the dorsomedial striatum impair serial spatial reversal learning in rats. Behav Brain Res 210:74–83.

Collins AGE, Frank MJ (2014) Opponent Actor Learning (OpAL): Modeling Interactive Effects of Striatal Dopamine on Reinforcement Learning and Choice Incentive. Psychol Rev 121:337–366.

Cox J, Witten IB (2019) Striatal circuits for reward learning and decision-making. Nat Rev Neurosci.

Cui G, Jun SB, Jin X, Pham MD, Vogel SS, Lovinger DM, Costa RM (2013) Concurrent activation of striatal direct and indirect pathways during action initiation. Nature 494:238–242.

Daw N (2009) Trial-by-trial data analysis using computational models. In: Decision Making, Affect, and Learning: Attention and Performance (Delgado MR, Phelps EA, Robbins TW, eds), pp 1–23: Oxford University Press.

Donahue CH, Liu M, Kreitzer AC (2018) Distinct value encoding in striatal direct and indirect pathways during adaptive learning. bioRxiv.

Dunovan K, Verstynen T (2016) Believer-Skeptic Meets Actor-Critic: Rethinking the Role of Basal Ganglia Pathways during Decision-Making and Reinforcement Learning. Frontiers in neuroscience 10:106.

Everitt BJ, Belin D, Economidou D, Pelloux Y, Dalley JW, Robbins TW (2008) Review. Neural mechanisms underlying the vulnerability to develop compulsive drug-seeking habits and addiction. Philos Trans R Soc Lond B Biol Sci 363:3125–3135.

Foerde K, Steinglass JE, Shohamy D, Walsh BT (2015) Neural mechanisms supporting maladaptive food choices in anorexia nervosa. Nature neuroscience 18:1571–1573.

Frank MJ, Seeberger LC, O’Reilly R C (2004) By carrot or by stick: cognitive reinforcement learning in parkinsonism. Science 306:1940–1943.

Franklin KBJ, Paxinos G (1997) The Mouse Brain in Stereotaxic Coordinates: Academic Press.

Freeze BS, Kravitz AV, Hammack N, Berke JD, Kreitzer AC (2013) Control of basal ganglia output by direct and indirect pathway projection neurons. The Journal of neuroscience : the official journal of the Society for Neuroscience 33:18531–18539.

Friedman A, Homma D, Gibb LG, Amemori K, Rubin SJ, Hood AS, Riad MH, Graybiel AM (2015) A Corticostriatal Path Targeting Striosomes Controls Decision-Making under Conflict. Cell 161:1320–1333.

Friedman A, Homma D, Bloem B, Gibb LG, Amemori KI, Hu D, Delcasso S, Truong TF, Yang J, Hood AS, Mikofalvy KA, Beck DW, Nguyen N, Nelson ED, Toro Arana SE, Vorder Bruegge RH, Goosens KA, Graybiel AM (2017) Chronic Stress Alters Striosome-Circuit Dynamics, Leading to Aberrant Decision-Making. Cell 171:1191–1205 e1128.

Geddes CE, Li H, Jin X (2018) Optogenetic Editing Reveals the Hierarchical Organization of Learned Action Sequences. Cell 174:32–43 e15.

Gerfen CR, Engber TM, Mahan LC, Susel Z, Chase TN, Monsma FJ Jr., Sibley DR (1990) D1 and D2 dopamine receptor-regulated gene expression of striatonigral and striatopallidal neurons. Science 250:1429–1432.

Gillan CM, Apergis-Schoute AM, Morein-Zamir S, Urcelay GP, Sule A, Fineberg NA, Sahakian BJ, Robbins TW (2015) Functional neuroimaging of avoidance habits in obsessive-compulsive disorder. Am J Psychiatry 172:284–293.

Hamid AA, Pettibone JR, Mabrouk OS, Hetrick VL, Schmidt R, Vander Weele CM, Kennedy RT, Aragona BJ, Berke JD (2016) Mesolimbic dopamine signals the value of work. Nature neuroscience 19:117–126.

Hikosaka O, Takikawa Y, Kawagoe R (2000) Role of the basal ganglia in the control of purposive saccadic eye movements. Physiological reviews 80:953–978.

Hikosaka O, Nakamura K, Nakahara H (2006) Basal ganglia orient eyes to reward. J Neurophysiol 95:567–584.

Howard WE, Marsh RE, Cole RE (1968) Food detection by deer mice using olfactory rather than visual cues. Anim Behav 16:13–17.

Humphries MD, Khamassi M, Gurney K (2012) Dopaminergic Control of the Exploration-Exploitation Trade-Off via the Basal Ganglia. Frontiers in neuroscience 6:9.

Isomura Y, Takekawa T, Harukuni R, Handa T, Aizawa H, Takada M, Fukai T (2013) Reward-modulated motor information in identified striatum neurons. The Journal of neuroscience : the official journal of the Society for Neuroscience 33:10209–10220.

Jin X, Tecuapetla F, Costa RM (2014) Basal ganglia subcircuits distinctively encode the parsing and concatenation of action sequences. Nature neuroscience 17:423–430.

Johnson C, Wilbrecht L (2011) Juvenile mice show greater flexibility in multiple choice reversal learning than adults. Dev Cogn Neurosci 1:540–551.

Johnson CM, Peckler H, Tai LH, Wilbrecht L (2016) Rule learning enhances structural plasticity of long-range axons in frontal cortex. Nature communications 7:10785.

Juavinett AL, Erlich JC, Churchland AK (2018) Decision-making behaviors: weighing ethology, complexity, and sensorimotor compatibility. Current Opinion in Neurobiology 49:42–50.

Kessler RM, Hutson PH, Herman BK, Potenza MN (2016) The neurobiological basis of binge-eating disorder. Neurosci Biobehav Rev 63:223–238.

Kim H, Sul JH, Huh N, Lee D, Jung MW (2009) Role of striatum in updating values of chosen actions. The Journal of neuroscience : the official journal of the Society for Neuroscience 29:14701–14712.

Kim HF, Amita H, Hikosaka O (2017) Indirect Pathway of Caudal Basal Ganglia for Rejection of Valueless Visual Objects. Neuron 94:920–930 e923.

Klaus A, Martins GJ, Paixao VB, Zhou P, Paninski L, Costa RM (2017) The Spatiotemporal Organization of the Striatum Encodes Action Space. Neuron 96:949.

Kravitz AV, Tye LD, Kreitzer AC (2012) Distinct roles for direct and indirect pathway striatal neurons in reinforcement. Nature neuroscience 15:816–818.

Kravitz AV, Freeze BS, Parker PR, Kay K, Thwin MT, Deisseroth K, Kreitzer AC (2010) Regulation of parkinsonian motor behaviours by optogenetic control of basal ganglia circuitry. Nature 466:622–626.

Kupferschmidt DA, Juczewski K, Cui G, Johnson KA, Lovinger DM (2017) Parallel, but Dissociable, Processing in Discrete Corticostriatal Inputs Encodes Skill Learning. Neuron 96:476–489 e475.

Lau B, Glimcher PW (2008) Value representations in the primate striatum during matching behavior. Neuron 58:451–463.

Lee E, Seo M, Dal Monte O, Averbeck BB (2015) Injection of a dopamine type 2 receptor antagonist into the dorsal striatum disrupts choices driven by previous outcomes, but not perceptual inference. The Journal of neuroscience : the official journal of the Society for Neuroscience 35:6298–6306.

Lucantonio F, Stalnaker TA, Shaham Y, Niv Y, Schoenbaum G (2012) The impact of orbitofrontal dysfunction on cocaine addiction. Nature neuroscience 15:358–366.

Mahler SV, Aston-Jones G (2018) CNO Evil? Considerations for the Use of DREADDs in Behavioral Neuroscience. Neuropsychopharmacology : official publication of the American College of Neuropsychopharmacology.

Mallet N, Ballion B, Le Moine C, Gonon F (2006) Cortical inputs and GABA interneurons imbalance projection neurons in the striatum of parkinsonian rats. The Journal of neuroscience : the official journal of the Society for Neuroscience 26:3875–3884.

Markowitz JE, Gillis WF, Beron CC, Neufeld SQ, Robertson K, Bhagat ND, Peterson RE, Peterson E, Hyun M, Linderman SW, Sabatini BL, Datta SR (2018) The Striatum Organizes 3D Behavior via Moment-to-Moment Action Selection. Cell 174:44–58 e17.

Matamales M, McGovern AE, Mi JD, Mazzone SB, Balleine BW, Bertran-Gonzalez J (2020) Local D2- to D1-neuron transmodulation updates goal-directed learning in the striatum. Science 367:549–555.

Mazzoni P, Hristova A, Krakauer JW (2007) Why don't we move faster? Parkinson's disease, movement vigor, and implicit motivation. The Journal of neuroscience : the official journal of the Society for Neuroscience 27:7105–7116.

Meng C, Zhou J, Papaneri A, Peddada T, Xu K, Cui G (2018) Spectrally Resolved Fiber Photometry for Multi-component Analysis of Brain Circuits. Neuron 98:707–717 e704.

Mink JW (1996) The basal ganglia: Focused selection and inhibition of competing motor programs. Prog Neurobiol 50:381–425.

Nonomura S, Nishizawa K, Sakai Y, Kawaguchi Y, Kato S, Uchigashima M, Watanabe M, Yamanaka K, Enomoto K, Chiken S, Sano H, Soma S, Yoshida J, Samejima K, Ogawa M, Kobayashi K, Nambu A, Isomura Y, Kimura M (2018) Monitoring and Updating of Action Selection for Goal-Directed Behavior through the Striatal Direct and Indirect Pathways. Neuron 99:1302–1314 e1305.

Ogasawara T, Nejime M, Takada M, Matsumoto M (2018) Primate Nigrostriatal Dopamine System Regulates Saccadic Response Inhibition. Neuron 100:1513–1526 e1514.

Parker JG, Marshall JD, Ahanonu B, Wu YW, Kim TH, Grewe BF, Zhang Y, Li JZ, Ding JB, Ehlers MD, Schnitzer MJ (2018) Diametric neural ensemble dynamics in parkinsonian and dyskinetic states. Nature 557:177–182.

Rescorla RA, Wagner AR (1972) A theory of Pavlovian conditioning: variations in the effectiveness of reinforcement and nonreinforcement. In: Classical Conditioning II: Current Research and Theory (Black A, Prokasy W, eds), pp 64–99. New York: Appleton Century Crofts.

Roseberry TK, Lee AM, Lalive AL, Wilbrecht L, Bonci A, Kreitzer AC (2016) Cell-Type-Specific Control of Brainstem Locomotor Circuits by Basal Ganglia. Cell 164:526–537.

Samejima K, Ueda Y, Doya K, Kimura M (2005) Representation of action-specific reward values in the striatum. Science 310:1337–1340.

Santacruz SR, Rich EL, Wallis JD, Carmena JM (2017) Caudate Microstimulation Increases Value of Specific Choices. Curr Biol 27:3375–3383 e3373.

Schmidt R, Leventhal DK, Mallet N, Chen F, Berke JD (2013) Canceling actions involves a race between basal ganglia pathways. Nature neuroscience 16:1118–1124.

Seo M, Lee E, Averbeck BB (2012) Action selection and action value in frontal-striatal circuits. Neuron 74:947–960.

Shan Q, Ge M, Christie MJ, Balleine BW (2014) The acquisition of goal-directed actions generates opposing plasticity in direct and indirect pathways in dorsomedial striatum. The Journal of neuroscience : the official journal of the Society for Neuroscience 34:9196–9201.

Shin JH, Kim D, Jung MW (2018) Differential coding of reward and movement information in the dorsomedial striatal direct and indirect pathways. Nature communications 9:404.

Surmeier DJ, Ding J, Day M, Wang ZF, Shen WX (2007) D1 and D2 dopamine-receptor modulation of striatal glutamatergic signaling in striatal medium spiny neurons. Trends in neurosciences 30:228–235.

Sutton RS, Barto AG (1998) Reinforcement learning : an introduction. Cambridge, Mass.: MIT Press.

Tai LH, Lee AM, Benavidez N, Bonci A, Wilbrecht L (2012) Transient stimulation of distinct subpopulations of striatal neurons mimics changes in action value. Nature neuroscience 15:1281–1289.

Thorn CA, Atallah H, Howe M, Graybiel AM (2010) Differential dynamics of activity changes in dorsolateral and dorsomedial striatal loops during learning. Neuron 66:781–795.

Vehtari A, Gelman A, Gabry J (2017) Practical Bayesian model evaluation using leave-one-out cross-validation and WAIC. Stat Comput 27:1413–1432.

Volkow ND, Morales M (2015) The Brain on Drugs: From Reward to Addiction. Cell 162:712–725.

Wang AY, Miura K, Uchida N (2013) The dorsomedial striatum encodes net expected return, critical for energizing performance vigor. Nature neuroscience 16:639-+.

Watanabe S (2010) Asymptotic Equivalence of Bayes Cross Validation and Widely Applicable Information Criterion in Singular Learning Theory. J Mach Learn Res 11:3571–3594.

Yin HH, Knowlton BJ, Balleine BW (2005a) Blockade of NMDA receptors in the dorsomedial striatum prevents action-outcome learning in instrumental conditioning. Eur J Neurosci 22:505–512.

Yin HH, Ostlund SB, Knowlton BJ, Balleine BW (2005b) The role of the dorsomedial striatum in instrumental conditioning. Eur J Neurosci 22:513–523.

Yin HH, Mulcare SP, Hilario MR, Clouse E, Holloway T, Davis MI, Hansson AC, Lovinger DM, Costa RM (2009) Dynamic reorganization of striatal circuits during the acquisition and consolidation of a skill. Nature neuroscience 12:333–341.

Yttri EA, Dudman JT (2016) Opponent and bidirectional control of movement velocity in the basal ganglia. Nature 533:402-+.

